# ProtoPure: Literature Mining and Analysis of Protein Purification Protocols at Proteome Scale

**DOI:** 10.64898/2026.10.26.762056

**Authors:** Ricardo Almada-Monter, Jose Martinez-Lomeli, Erika Garay, Joaquίn Trevilla-Santa Ana, Amanda Spencer, Adriana Siordia, Param Chordiya, Sofίa Luna-Armijo, Javier Espinoza-Herrera, Nathaniel Roethler, Sarah Marta Veskimägi, Akshay Uppal, Melina Shamshoum, Adrian Jinich

**Affiliations:** Department of Chemistry and Biochemistry, University of California San Diego, La Jolla, California 92093, United States; Department of Molecular, Cell and Systems Biology, University of California, Riverside, California, United States; Departamento de Matemáticas, Facultad de Ciencias, UNAM, Ciudad Universitaria, Circuito Exterior, Cd. de México, C.P., 04510, México; Skaggs School of Pharmacy and Pharmaceutical Sciences, University of California San Diego, La Jolla, California 92093, United States; Department of Electrical & Computer Engineering, University of California San Diego, La Jolla, California 92093, United States; Instituto Tecnológico de Tijuana, Tijuana, Baja California, México; Universidad Autónoma de Baja California, Tijuana, Baja California, México; Halıcıoğlu School of Data Science and Computing, University of California San Diego, La Jolla, California, 92093, United States

**Keywords:** protein expression, literature mining, AI for Science, knowledge discovery, protocol standardization, semantic clustering, AI for protein design, literature-derived datasets

## Abstract

Protein purification is essential for characterizing protein structure and function, yet finding the optimal conditions is a laborious task that requires extensive experimentation. Researchers usually survey the literature to identify starting conditions, which are optimized for the protein of interest. To facilitate this task, we introduce ProtoPure, an LLM framework to extract and analyze protein purification conditions at the protein family or proteome scale. We evaluated ProtoPure against a curated ground-truth dataset of *∼*100 publications across multiple metrics (BLEU, METEOR, BERTScore, and LLM-based scoring), achieving a mean LLM-based score of 7.59 out of 10 across all extracted parameters. We then applied ProtoPure to construct a dataset spanning three distinct protein families and three proteomes (*Escherichia coli*, *Mycobacterium tuberculosis* and *Thermus thermophilus*), analyzing over 10,000 publications and extracting purification conditions for more than 12,000 protein records.

The analysis of the resulting dataset revealed a strong convergence toward a limited set of canonical protein purification workflows, characterized by widespread heterologous expression in the BL21(DE3) expression system alongside more flexible, context-dependent variation in plasmid selection, growth media, desalting strategies, and elution buffer composition.

We make the protein purification records reconstructed in this study publicly available to the research community through the ProtoPure database. The platform allows researchers to explore purification conditions reported for proteins of interest, compare experimental strategies across related proteins, families, or organisms, and trace individual records back to their source publications.

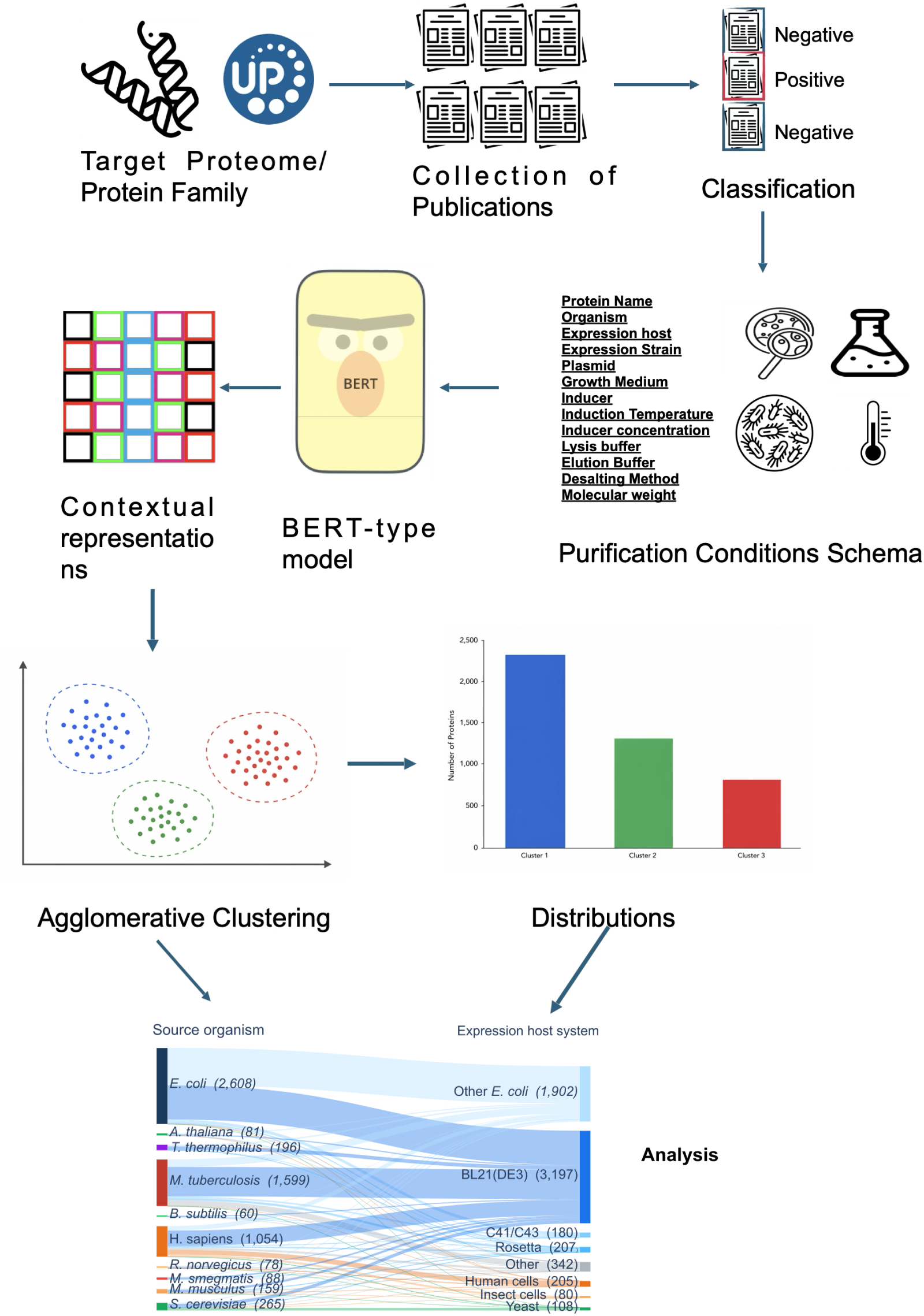
Graphical Abstract.

**The Bigger Picture:** Protein purification is a fundamental step for analyzing individual proteins and protein complexes and identifying interactions with other proteins, DNA, or RNA. A variety of protein purification strategies exist to address desired scale, throughput, and downstream applications. However, selecting an effective experimental procedure remains a largely empirical process that a quick literature review can help guide. Researchers typically survey on the order of a dozen publications for proteins with similar biochemical properties, assessing conditions on a case-by-case basis, leaving no systematic framework for comparison and decision-making at the scale of entire protein families. As protein engineering and design evolve, identifying effective purification conditions remains critical for obtaining sufficient quantities of properly folded and functional proteins.

To address this gap, we developed ProtoPure, an LLM-based workflow and database that automates the extraction, standardization and analysis of protein purification conditions from peer-reviewed literature at scale. Given a user-defined UniProt table—enabling flexible selection of proteins by family, organism, or proteome—ProtoPure retrieves associated publications, classifies them based on the presence of purification protocols, and extracts a standardized set of 12 purification parameters (e.g., buffer composition, elution conditions). By transforming dispersed experimental descriptions into a structured resource, ProtoPure makes it possible to compare purification strategies across thousands of proteins. The resulting dataset revealed quantitative convergence toward a limited set of recombinant expression practices, with BL21(DE3)-based systems dominating protein production and recurrent combinations of expression host, inducer, growth medium, and plasmid occurring substantially more frequently than expected by chance. In contrast, downstream purification parameters exhibited greater diversity, revealing a workflow architecture that combines standardized expression infrastructure with more flexible biochemical components.

**Highlights:**

- ProtoPure extracts protein purification protocols from scientific literature.
- ProtoPure converts unstructured protocols into a searchable biological resource.
- Purification workflows converge toward a limited set of common architectures.
- Expression systems are highly standardized, while downstream steps remain flexible.
- Literature-derived protocols enable data-driven experimental design and optimization.
- The resulting resource enables evidence-based comparison of purification conditions.

## INTRODUCTION

Protein purification is a cornerstone of experimental biochemistry, aimed at isolating a target protein from a heterogeneous cellular environment in order to study its structure and function. Proteins sharing similar physical or biochemical properties often follow comparable purification protocols, allowing researchers to adapt well-established methods from the literature to purify novel proteins.

As protein design becomes more prominent, establishing a protein purification protocol is essential to obtain sufficient quantities of protein for characterization while maintaining correct folding, stability, and functional integrity. However, despite rapid advances in computational protein design, there is currently no large-scale resource that systematically links proteins to experimentally validated purification conditions, leaving protocol selection largely dependent on manual literature searches and empirical optimization.

Recent advances in Large Language Models (LLMs) have transformed how scientific research is conducted, enabling the automation of many tasks that would otherwise demand a substantial amount of time and expertise. In chemistry and biological applications, LLMs have gained traction for their generative capabilities, such as computer-aided synthesis planning and reaction prediction^1,2^, molecular property prediction^3–5^ and virtual screening^6–8^.

In the field of experimental biochemistry, extracting protocols from published literature at scale requires extensive manual curation and substantial domain expertise to interpret experimental designs, identify relevant variables, and standardize measurements across studies. This process is time-consuming and susceptible to human bias and inconsistency, limiting scalability and reproducibility.

LLMs have been increasingly used to extract and standardize information across scientific domains, including biological metadata and ontology terms, ecological and gene-related data, and biomedical concepts^9–14^. Similar approaches have also been applied in chemistry and materials science to recover synthesis conditions for metal–organic frameworks and material properties such as electronic band gaps from peer-reviewed literature^15–21^.

Only limited prior work has explored large-scale extraction of protein purification conditions from peer-reviewed literature using LLMs^22^. This prior work focused primarily on PDB-associated publications, leaving proteins studied for other purposes, such as enzymology, underrepresented. The limited adoption of such approaches likely reflects the diversity of purification protocols, variability in reporting formats, and reliance on specialized terminology.

In this study, we investigated protein purification strategies across entire protein families and proteomes. ProtoPure automatically retrieves publications associated with UniProt entries, identifies publications that report purification protocols, extracts the method section of the publication, reconstructs purification conditions according to a standardized schema, and groups related protocols using contextual biological embeddings. Unlike previous PDB-centered approaches, ProtoPure enables systematic analysis across arbitrary UniProt-derived protein families and complete proteomes.

This approach allows us to identify recurring purification strategies, family-specific trends, and shared experimental patterns across multiple proteomes that are difficult to capture using manual curation or rule-based extraction methods.

## RESULTS

### ProtoPure recovers purification conditions with high semantic fidelity across twelve parameters

To ensure that the large-scale analyses were based on reliable data, we evaluated each stage of ProtoPure, from publication classification to purification-condition extraction. To perform this evaluation, we employed datasets from two enzyme families, SAM-dependent methyltransferases (SAM-MTases) and azoreductases. Details regarding the retrieval and manual curation of these datasets are provided in the Methods section.

ProtoPure begins with a UniProt-derived table of proteins of interest; however, not all associated publications report protein purification conditions. We therefore evaluated publication-level classification using the SAM-MTase dataset, followed by methods-section extraction and purification-condition extraction against manually curated records from both the SAM-MTase and azoreductase datasets. Publication classification was implemented using LlamaParse, whereas methods-section extraction and purification-condition reconstruction were performed using models accessed through the OpenAI API.

Rather than optimizing for verbatim retrieval, we prioritized the extraction of information that was semantically and contextually equivalent to the manually curated ground truth. Accordingly, we evaluated ProtoPure using semantic similarity metrics, including BERTScore^23^, together with an LLM-as-a-judge evaluation. LLM-as-a-judge approaches, in which a language model evaluates the quality or agreement of model-generated outputs against predefined criteria or reference information, have been increasingly used for evaluating outputs in tasks where conventional lexical metrics may not adequately capture semantic equivalence^24–28^. In our case, the model was provided with both the manually curated ground-truth record and the corresponding ProtoPure-extracted information and asked to assess their semantic agreement on a scale from 0 to 10. A score of 10 indicated that the extracted and ground-truth information were equivalent in meaning, whereas a score of 0 indicated unrelated or contradictory information. Representative LLM-based similarity scores and their corresponding evaluation rationales are shown in **Table 2** to showcase why we use this metric as the main evaluator of the pipeline. For example, ProtoPure extracted Homo sapiens where the manually curated record contained ‘Human’ (**Table 2**). The LLM-based evaluator correctly assigned this pair a score of 10 because the terms are semantically equivalent despite their lexical differences.

**Table 1.** Protein purification schema used by ProtoPure.

| Field | Description |
| --- | --- |
| Protein name | Name of the purified target protein |
| Organism source | Organism from which the protein originates |
| Organism Strain | Organism or system used for recombinant expression |
| Expression strain | Specific strain used for protein expression |
| Plasmid | Expression vector used for protein production |
| Growth medium | Culture medium used during expression |
| Inducer | Chemical or condition used to induce protein expression |
| Induction temperature | Temperature used during induction |
| Lysis buffer | Buffer used for protein extraction following cell disruption |
| Elution buffer | Buffer or condition used to elute the purified protein |
| Desalting method | Method used for buffer exchange or desalting |
| Molecular weight | Reported molecular weight of the purified protein |

The table describes the schema used by ProtoPure to extract and standardize protein purification conditions from scientific publications.

**Table 2.**
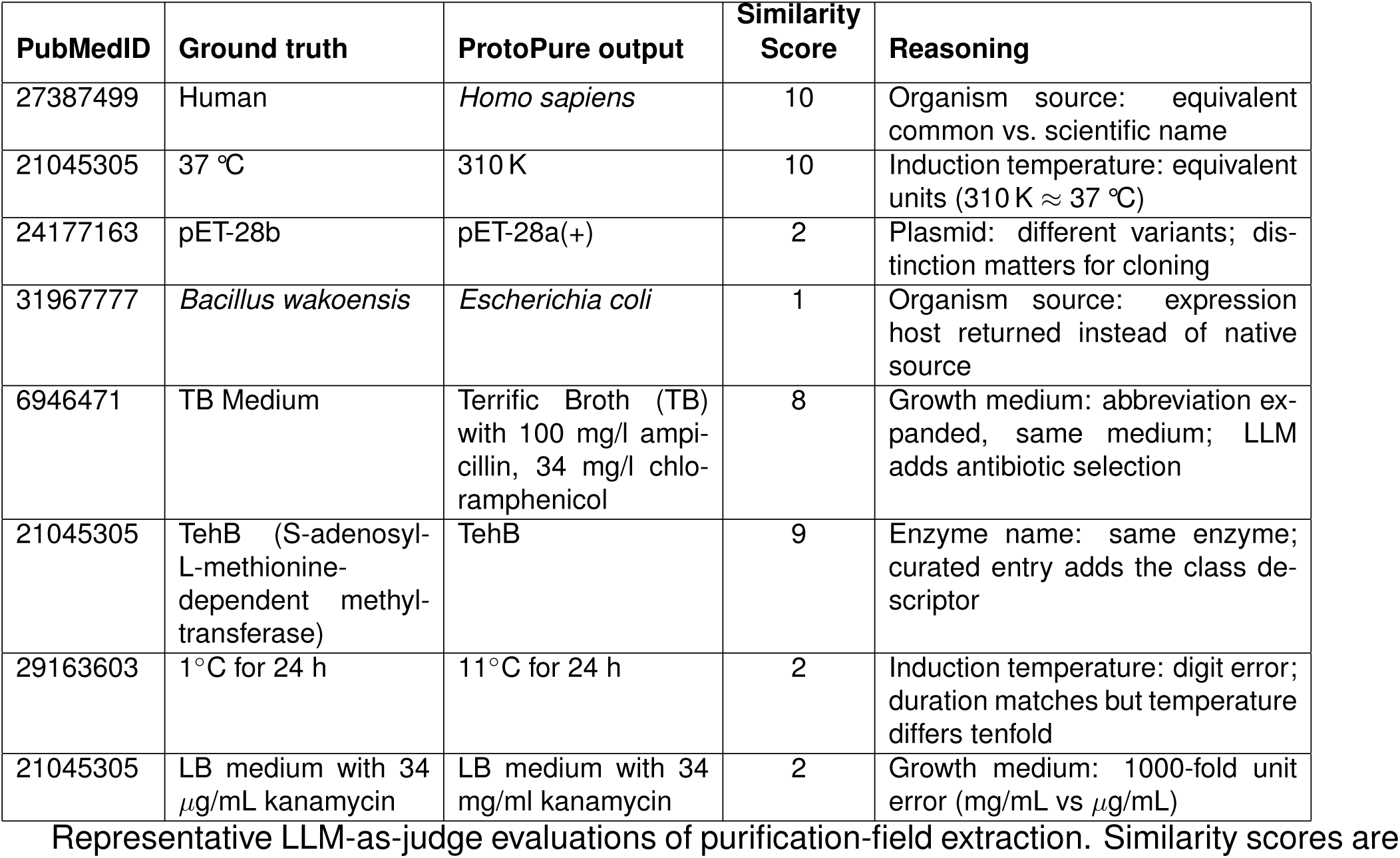
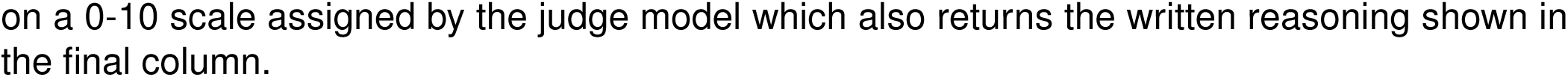
Examples of Evaluation Scores with LLMs.

| PubMedID | Ground truth | ProtoPure output | Similarity Score | Reasoning |
| --- | --- | --- | --- | --- |
| 27387499 | Human | *Homo sapiens* | 10 | Organism source: equivalent common vs. scientific name |
| 21045305 | 37 °C | 310 K | 10 | Induction temperature: equivalent units (310 K ≈ 37 °C) |
| 24177163 | pET-28b | pET-28a(+) | 2 | Plasmid: different variants; distinction matters for cloning |
| 31967777 | *Bacillus wakoensis* | *Escherichia coli* | 1 | Organism source: expression host returned instead of native source |
| 6946471 | TB Medium | Terrific Broth (TB) with 100 mg/l ampicillin, 34 mg/l chloramphenicol | 8 | Growth medium: abbreviation expanded, same medium; LLM adds antibiotic selection |
| 21045305 | TehB (S-adenosyl-L-methionine-dependent methyltransferase) | TehB | 9 | Enzyme name: same enzyme; curated entry adds the class descriptor |
| 29163603 | 1°C for 24 h | 11°C for 24 h | 2 | Induction temperature: digit error; duration matches but temperature differs tenfold |
| 21045305 | LB medium with 34 µg/mL kanamycin | LB medium with 34 mg/ml kanamycin | 2 | Growth medium: 1000-fold unit error (mg/mL vs µg/mL) |

Representative LLM-as-judge evaluations of purification-field extraction. Similarity scores are

On the SAM-MTase classification benchmark (**Fig. 1A**), ProtoPure achieved an F1 score of 77.7% and an accuracy of 85.3%. The classifier showed high recall (88.2%) but lower precision (69.4%), correctly identifying 75 of the 85 publications containing protein purification information while misclassifying 33 publications without purification information as positive. This indicates that the classifier favors sensitivity over precision, capturing the majority of relevant publications at the cost of retaining some false positives. This behavior is suitable for the downstream extraction workflow, where false-positive publications can subsequently yield no purification records, whereas false negatives would exclude potentially relevant purification information from further analysis.

**Figure 1.**
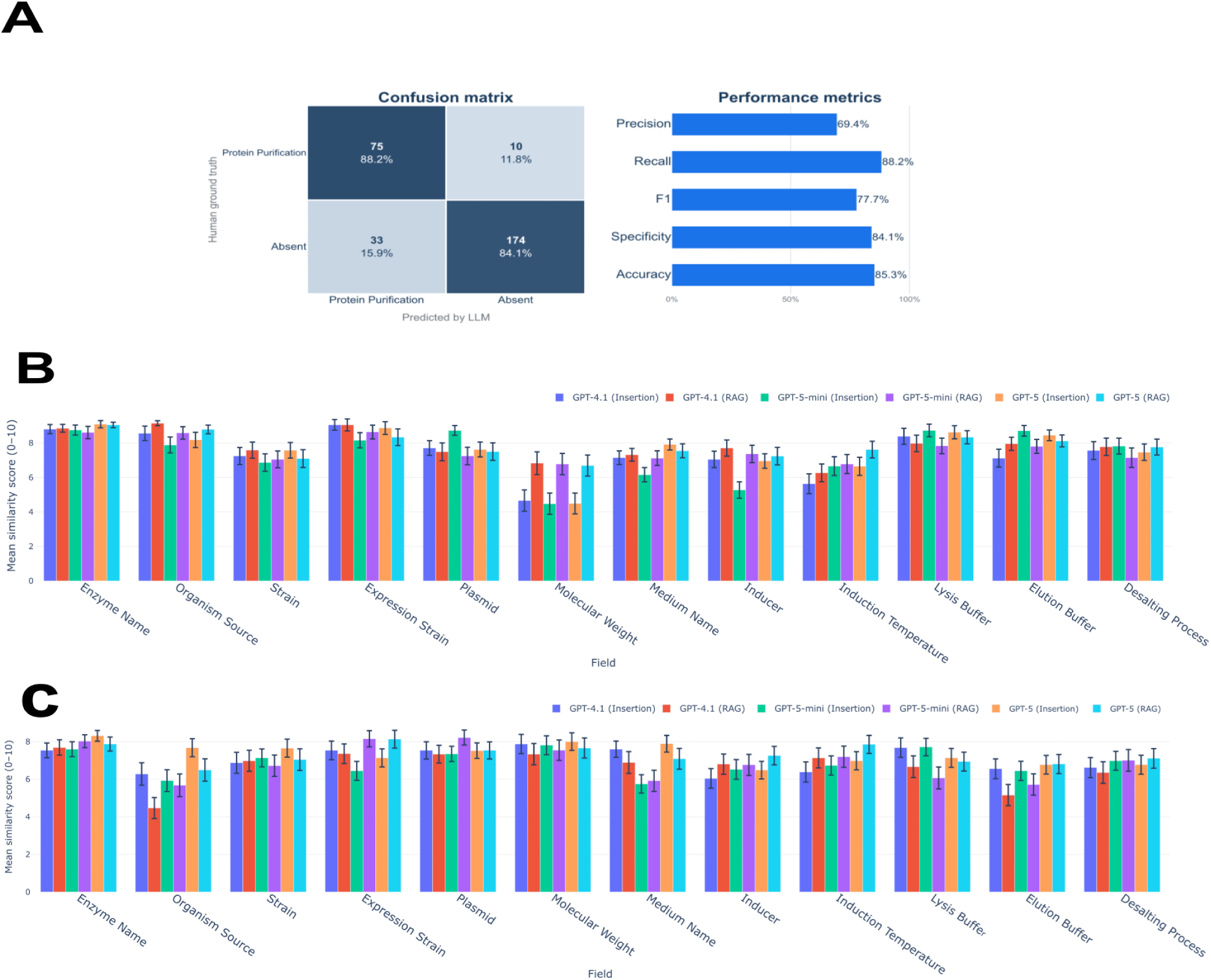
ProtoPure accurately reconstructs protein purification protocols from the scientific literature. Agreement between ProtoPure-extracted purification conditions and manually curated ground-truth records across publication classification and individual fields of the protein purification schema. Classification of publications according to the presence of protein purification conditions was evaluated using precision, recall, and F1 score. Agreement between extracted and manually curated purification records was evaluated using an LLM-as-a-judge on a scale from 0 to 10, where 10 indicates equivalent information and 0 indicates unrelated or contradictory information. Bars represent the mean similarity score for each purification field, and error bars indicate the standard error of the mean (SEM) across records. (**A**) Classification performance on the SAM-MTase benchmark dataset. Classification was not evaluated for the azoreductase dataset because this dataset was manually curated to include publications containing protein purification conditions. (**B**) Field-level extraction performance on the azoreductase benchmark dataset. (**C**) Field-level extraction performance on the SAM-MTase benchmark dataset.

Performance of the methods-section extraction step was evaluated using conventional natural language processing metrics, including BLEU, METEOR, and ROUGE (**Fig. S1**). These metrics were used for this stage because methods sections are extracted verbatim from the source publications.

For purification-condition extraction (**Fig. 1B–C**), ProtoPure consistently recovered purification information with high semantic fidelity across evaluated fields and models. Mean similarity scores (LLM as a Judge metric) were predominantly above 7, with an overall mean of 7.59 (95% CI [7.48–7.70]). The highest agreement was observed for enzyme name (8.86, 95% CI [8.66– 9.07]) and organism source (8.51, 95% CI [8.21–8.81]), whereas molecular weight consistently showed the lowest agreement across models (5.63, 95% CI [5.12–6.14]).

These observations suggest that certain purification attributes are more challenging to extract than others. Variation in performance across fields may reflect both how consistently an attribute is reported and how readily it can be identified within a publication. Fields such as protein name and organism source are typically reported explicitly and unambiguously and were recorded by curators in 100% and 98% of protein records, respectively. In contrast, attributes such as molecular weight, induction temperature, and inducer were reported less consistently, appearing in 35%, 57%, and 69% of curated records, respectively. These differences in reporting completeness may contribute to the observed variation in extraction performance across purification fields. Overall, all models were efficient in extracting information showing minimal variations in extraction performance.

Across all tested models, performance was largely consistent, with differences between models generally falling within overlapping confidence intervals. GPT-5 showed a marginal overall advantage (mean similarity 7.66 vs. 7.41 and 7.35 for GPT-4.1 and GPT-5-mini, respectively, on the azoreductase benchmark). Nevertheless, no model consistently outperformed the others across purification fields, with GPT-4.1 and GPT-5-mini achieving comparable or higher performance for individual attributes.

As a complementary measure of semantic similarity, we also evaluated the extracted purification conditions using BERTScore^23^ (**Fig. S2**). BERTScore values were generally high across purification fields and were broadly consistent across the evaluated language models and extraction strategies. Differences were more apparent between purification fields than between model configurations, further supporting that ProtoPure reliably captures the semantic content of manually curated purification records.

Together, these results demonstrate that ProtoPure reliably recovers protein purification conditions from the literature, enabling their systematic analysis at scale.

### ProtoPure enables proteome and protein family-scale reconstruction of protein purification conditions

Once we established that ProtoPure could reliably reconstruct purification conditions across all stages of the workflow, we applied it to a large-scale analysis spanning three protein families: short-chain dehydrogenases/reductases (SDRs), small GTP-binding domain proteins (SgtPB), the heat shock protein 70 (HSP-70); family and the complete proteomes of three organisms: *Escherichia coli*, *Mycobacterium tuberculosis*, and *Thermus thermophilus*. These protein families and proteomes were selected because they represent well-studied and well-characterized biological systems with substantial representation in the scientific literature. In total, 10,908 publications were screened for evidence of protein purification, of which 4,582 (42.0%) were classified as reporting explicit purification conditions. From these, ProtoPure reconstructed purification conditions for 12,179 protein records. Among publications associated with *E. coli*, *M. tuberculosis*, and *T. thermophilus*, 38.9%, 76.3%, and 74.6%, respectively, contained protein purification methods. Similarly, 33.1%, 50.5%, and 9.0% of publications associated with SgtPB, SDRs, and HSP-70 proteins, respectively, reported protein purification methods (**Fig. 2**).

**Figure 2.**
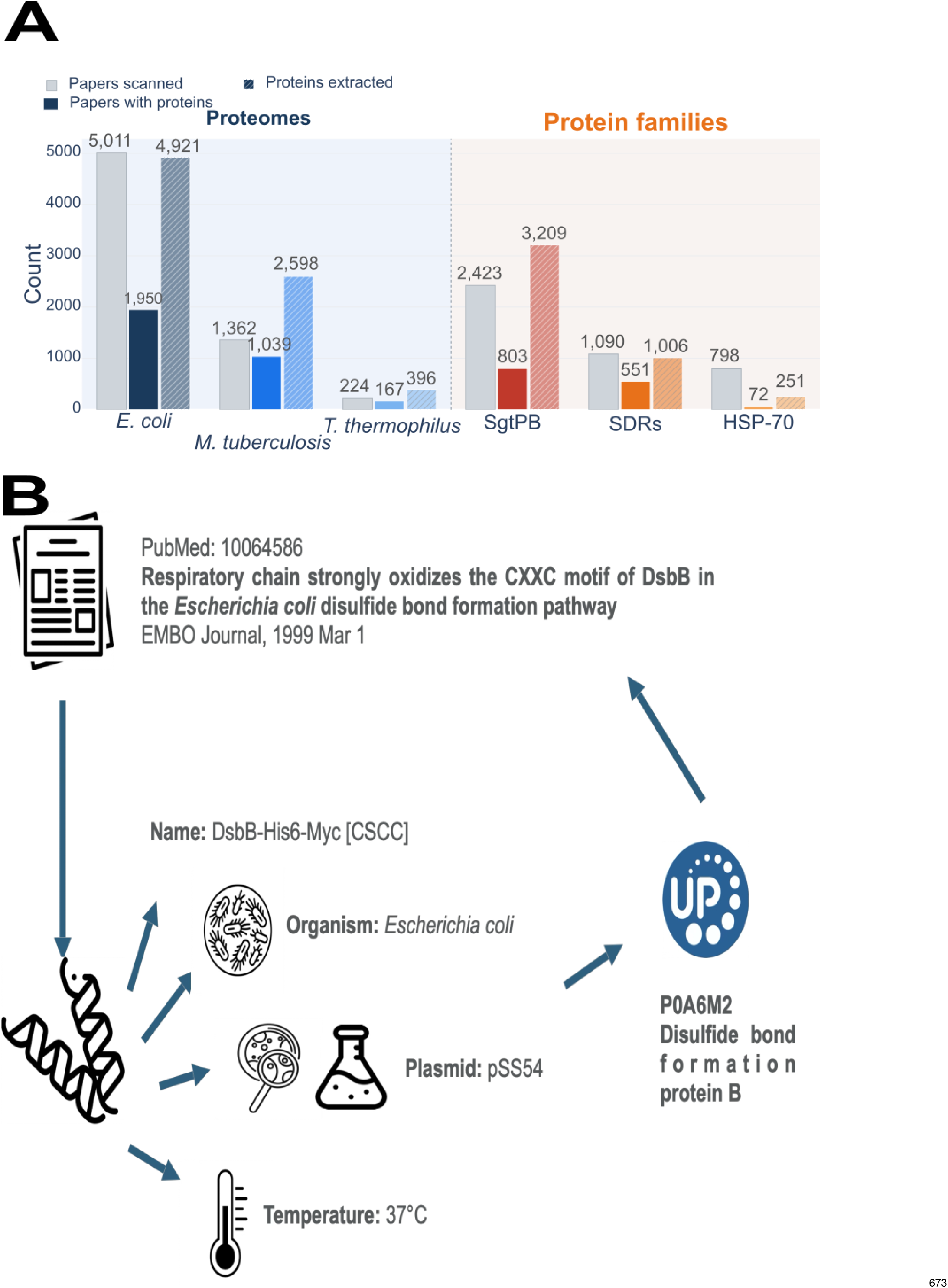
ProtoPure enables large-scale reconstruction and exploration of protein purification conditions. (**A**) Scale of the datasets analyzed across three complete proteomes and three protein families. The proteome datasets comprise *Escherichia coli* (*E. coli*), *Mycobacterium tuberculosis* (*M. tuberculosis*), and *Thermus thermophilus* (*T. thermophilus*), while the protein-family datasets comprise small GTP-binding proteins (SgtPB), short-chain dehydrogenases/reductases (SDRs), and heat shock protein 70 (HSP-70). Publications associated with UniProt/Swiss-Prot entries were retrieved and screened for the presence of protein purification information, followed by extraction and standardization of the reported purification conditions. The numbers shown indicate the publications screened, publications classified as containing protein purification protocols, and protein records reconstructed for each dataset. (**B**) Representative entry from the ProtoPure resource showing how an extracted purification record retains its connection to the source publication and corresponding protein/UniProt entry while organizing the reported experimental conditions into the standardized purification schema.

To gain insight into the annotations of proteins for which purification information was available, each protein and its corresponding purification data were mapped to UniProtKB/Swiss-Prot accessions. Of the 12,179 purification records, 6,621 (54.4%) were directly mapped. For these mapped records, we retrieved additional protein annotations from UniProt, including cellular localization, Enzyme Commission (EC) number, UniProt keywords, reported cofactors, and protein family annotations.

The distributions of these protein annotations across the mapped dataset are shown in **Fig. 3**. Among the 4,493 proteins with a confidently assigned cellular localization, nearly half were cytoplasmic (47.3%). Membrane-associated proteins represented the second-largest group (26.9%) when cell, inner, and outer membrane annotations were combined, whereas secreted (6.5%), periplasmic (4.6%), and organellar proteins (endoplasmic reticulum, 3.7%; nucleus, 3.1%; mitochondrion, 2.6%) were less frequently represented. Among the 4,422 proteins with EC annotations, hydrolases (36.0%), transferases (27.0%), and oxidoreductases (21.8%) predominated, together accounting for 84.8% of EC-annotated proteins, while lyases, isomerases, ligases, and translocases comprised the remaining classes. The most frequently represented functional keywords were metal-binding (29.7%) and nucleotide-binding (28.8%). Finally, among the 2,088 proteins with annotated cofactors, divalent metal ions predominated, particularly Mg^2+^ (51.4%), Zn^2+^ (16.5%), and Mn^2+^ (14.9%). Organic and complex cofactors, including FAD, [4Fe-4S] clusters, and pyridoxal 5*^′^*-phosphate, each accounted for approximately 5% of cofactor annotations. Together, these annotations demonstrate that the mapped ProtoPure records span proteins with diverse cellular, enzymatic, and functional properties, providing a biologically diverse dataset for investigating common and context-dependent purification strategies.

**Figure 3.**
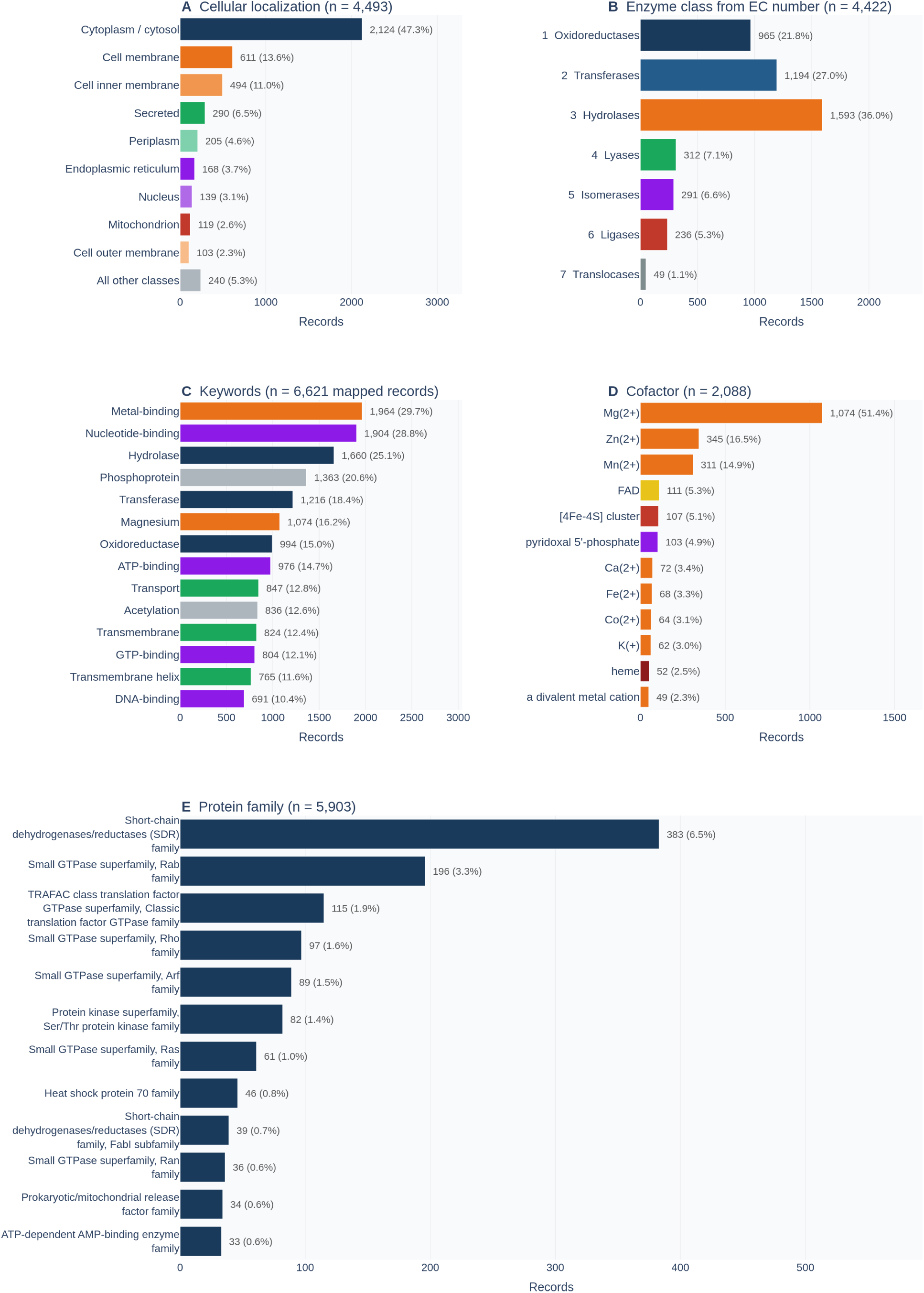
Biological and functional characteristics of proteins represented in the ProtoPure database. (**A**) Distribution of cellular localization annotations among proteins with an assigned UniProt cellular localization (*n* = 4, 493). (**B**) Distribution of enzyme classes based on Enzyme Commission (EC) numbers among EC-annotated proteins (*n* = 4, 422). (**C**) Most frequently occurring UniProt keywords among mapped purification records (*n* = 6, 621). (**D**) Distribution of annotated cofactors among proteins with UniProt cofactor annotations (*n* = 2, 088). (**E**) Most frequently represented UniProt protein families among proteins with family annotations (*n* = 5, 903).

To facilitate reuse of the extracted information, all reconstructed purification conditions are made available through the ProtoPure platform. Users can query proteins by UniProt accession, protein family, organism, or proteome, inspect the original publication associated with each protocol, and download standardized purification annotations for downstream analyses (**Fig. 2B**). Having established a large-scale resource of protein purification records, we next used the aggregated dataset to investigate broader patterns in purification conditions.

### Large-scale reconstruction reveals heterogeneous reporting completeness

We first investigated how consistently individual purification parameters were reported across the reconstructed dataset. Analysis of the extracted purification conditions revealed substantial variability in reporting practices across experimental fields (See **Fig. 4A**). Protein name and organism source were reported for the vast majority of entries (99.3% and 80.7%, respectively). In contrast, several experimentally critical purification parameters were reported considerably less frequently. Expression host, plasmid, and growth medium were present in 62.6–67.9% of protein records, whereas induction conditions, lysis buffers, elution buffers, and desalting procedures were only explicitly reported in 42.2–45.9% of cases. Molecular weight was the least reported field overall, appearing in only 4.1% of extracted purification records.

**Figure 4.**
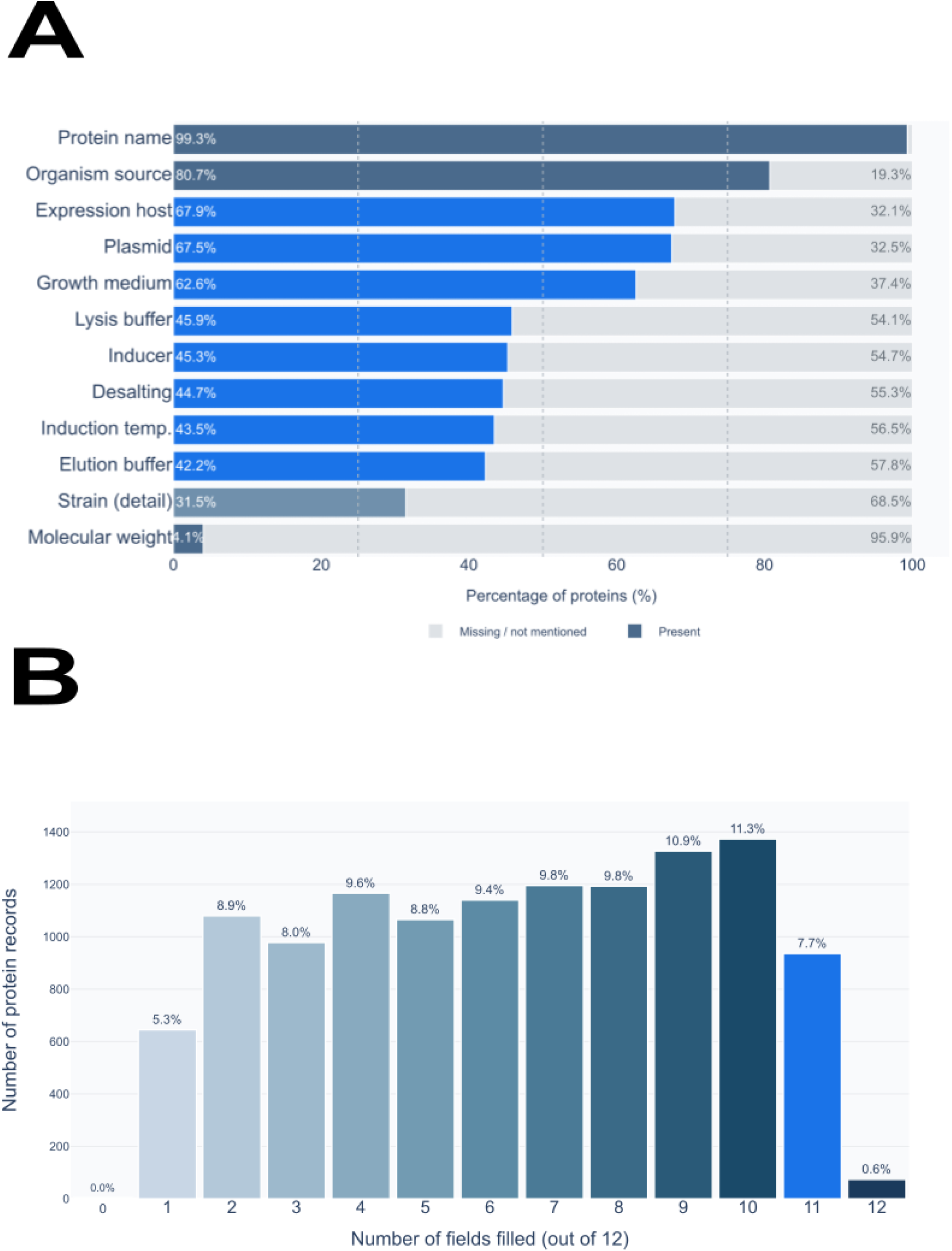
Protein purification parameters are incompletely and unevenly reported across the literature. (**A**) Reporting completeness across the individual fields of the protein purification schema. Bars indicate the percentage of reconstructed protein purification records in which each experimental parameter was reported. (**B**) Distribution of the number of purification fields reported per reconstructed protein record, illustrating variation in the overall completeness of purification protocols across the literature.

The overall completeness distribution showed a median of seven populated schema fields per protein record, with 59.4% of records containing six or more. Most publications provide substantial experimental detail despite not documenting fully exhaustive purification workflows. Complete reporting across all twelve fields was rare, occurring in only 0.6% of records (74 of 12,179). These results highlight substantial differences in the extent to which individual experimental parameters are documented in the literature.

To systematically compare the reported purification conditions across studies, we next transformed the extracted text into contextual representations. This allowed diverse descriptions of the same or related experimental conditions to be analyzed as standardized groups, enabling systematic comparison of their frequency, diversity, and associations across proteins.

To identify an appropriate representation for grouping semantically similar purification conditions, we evaluated multiple biomedical and general-purpose text encoders across a range of clustering parameters. We looked for clusters that maximized within-cluster similarity and between-cluster separation, as assessed using the Silhouette score and Davies–Bouldin index, while retaining sufficient cluster diversity to represent distinct purification conditions. The complete grid-search results are provided in the Supplemental Information (**Figs. S3–S13**). Examples of similar protocols expressed in different formats are shown in **Table 3**.

**Table 3.** Examples of members of an elution-buffer cluster (Cluster 1)

| PMID | Extracted value (verbatim) |
| --- | --- |
| 38740275 | Ni-NTA elution buffer (500 mM NaCl, 50 mM sodium phosphate, 250 mM imidazole, pH 8.0) |
| 24731824 | 50 mM NaH<sub>2</sub>PO<sub>4</sub>, 300 mM NaCl, 250 mM imidazole, pH 8.0 (Ni-NTA elution) |
| 30560095<sup>a</sup> | 50 mM sodium phosphate buffer (pH 8.0), 300 mM NaCl, 300 mM imidazole (Ni-NTA) |
| 18423586 | 50 mM sodium phosphate buffer, pH 8.0, 300 mM NaCl, 250 mM imidazole (Ni<sup>2+</sup>-NTA elution) |

<sup>a</sup>Contributes six rows to the cluster, one per protein reported in that publication; the nine cluster members comprise four distinct wordings. 756

### Purification protocols contain standardized and flexible components

We next investigated whether purification practices converged toward a limited set of commonly used experimental conditions or remained highly diverse across proteins. Despite sourcing proteins from 5 kingdoms of life and 462 distinct organisms, 87.7% of all purification experiments were conducted in *Escherichia coli*, with 51.0% using a single strain, BL21(DE3). Of the 63.5% of proteins expressed heterologously (i.e., in a host from a different organism than their source organism), 80.6% were still directed to *E. coli*. This collapse is most pronounced for biologically distant proteins: plant proteins were expressed in *E. coli* at a rate of 96.9%, and protist proteins at 98.0%, despite the substantial evolutionary distance and fundamental differences in post-translational machinery. Together, these findings provide an empirical, quantitative view of a well-established practice in recombinant protein production: the widespread use of *E. coli* as an expression host. Rather than simply confirming its known prevalence, our analysis quantifies the extent of this convergence across diverse protein source organisms, showing that expression is concentrated within a limited set of eight host systems, with *E. coli* BL21(DE3) alone accounting for the largest fraction of experiments (**Fig. 5**).

**Figure 5.**
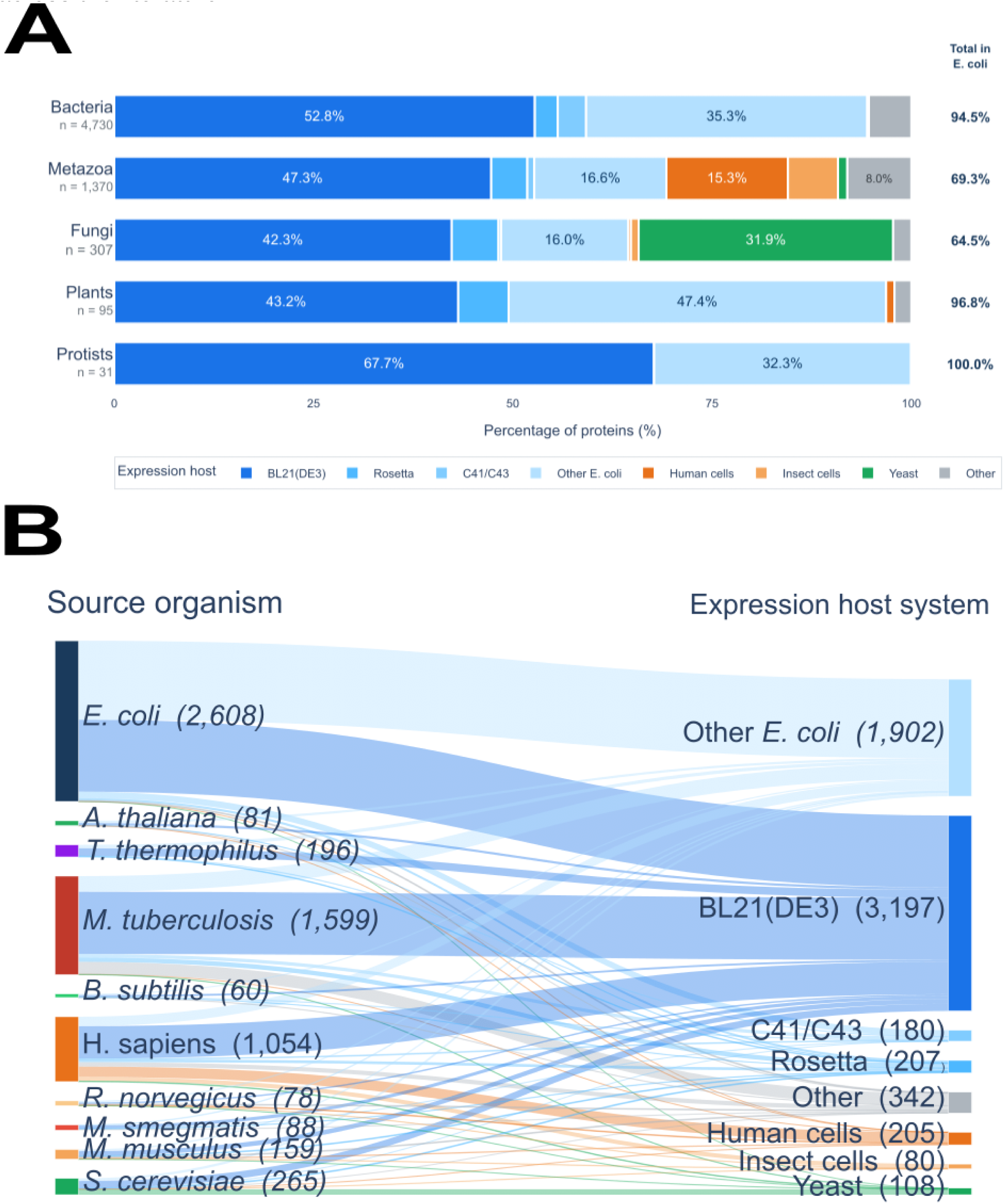
Protein purification workflows converge toward a limited set of expression systems. (**A**) Distribution of expression systems used for recombinant production of proteins originating from different taxonomic groups. Bars show the relative frequency of expression hosts within each protein-source group. (**B**) Sankey diagram connecting protein source organisms to the expression systems used for recombinant protein production. Flow width is proportional to the number of reconstructed protein purification records associated with each source-organism–expression-system combination, illustrating the convergence of diverse protein sources toward a limited number of commonly used expression systems.

Cluster analysis (**Fig. 6**) revealed substantial differences in the distribution of purification conditions across fields. Some fields were dominated by a small number of highly prevalent clusters, whereas others exhibited greater diversity, with observations distributed across many clusters. To quantify these differences, we calculated the Shannon entropy of each field from the relative abundance of its clusters and converted it to an effective number of clusters. This metric accounts for both the number of clusters and their relative frequencies, providing a measure of the effective diversity of purification conditions within each field.

**Figure 6.**
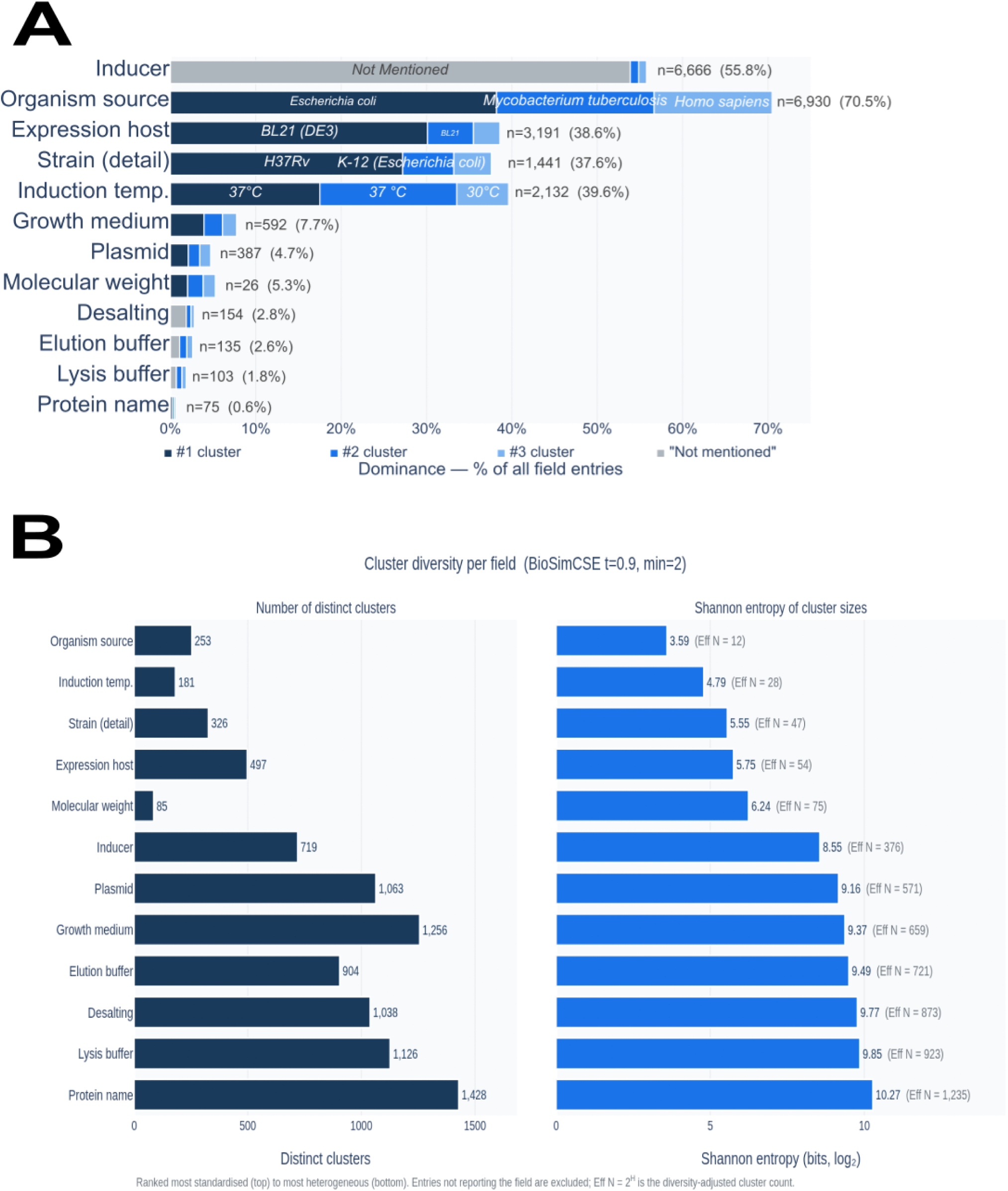
Protein purification parameters vary widely in standardization and experimental diversity. (A) Distribution of the most prevalent semantic clusters identified for each protein purification parameter. Bars indicate the relative abundance of the dominant clusters, illustrating the extent to which individual parameters are concentrated around a limited number of commonly reported experimental conditions. (**B**) Diversity of semantic clusters across protein purification parameters. Raw cluster counts represent the total number of distinct semantic clusters identified for each parameter, whereas the effective number of clusters, derived from Shannon entropy, accounts for differences in cluster abundance and reflects the number of substantially represented experimental alternatives. Higher Shannon entropy corresponds to a more even distribution of observations across clusters and therefore greater effective diversity, whereas lower entropy indicates that observations are concentrated within a smaller number of dominant clusters, reflecting greater standardization.

Expression-related parameters showed particularly strong convergence. Among entries reporting an expression host, BL21(DE3) represented the dominant condition, while IPTG accounted for 86.3% of entries reporting an inducer. Expression host exhibited an effective diversity of only 54 clusters, consistent with the concentration of recombinant expression around a relatively restricted set of host systems. Although semantic clustering resolved inducer descriptions into a comparatively large number of clusters, inspection of the most populated clusters revealed that the top clusters predominantly represented variants of IPTG that differed in associated information such as concentration, duration, or textual description. Thus, the apparent cluster diversity of this field largely reflects methodological fragmentation of a common underlying experimental condition rather than diversity in inducer identity. Induction temperature similarly showed substantial convergence, with 53.6% of reported values falling between 30*^◦^*C and 37*^◦^*C and an effective diversity of only 28 clusters. In contrast, plasmid selection, lysis buffer composition, and elution conditions exhibited substantially greater diversity, with effective diversities of 571, 923, and 721 clusters, respectively, and their largest individual clusters accounting for only 3.6%, 1.0%, and 1.4% of reported entries.

Together, these findings reveal that the diversity of proteins studied across the literature does not translate directly into equivalent diversity in experimental practice. Instead, proteins from widely different biological origins converge onto a relatively restricted recombinant expression infrastructure, while downstream purification conditions show more diversity in their respective fields.

### BL21(DE3), IPTG, LB, and pET vectors dominate protein expression workflows

We next investigated whether the observed convergence extended beyond individual purification parameters, asking whether specific experimental conditions consistently co-occurred to define recurrent purification workflows. Analysis of the four fields that jointly define a recombinant expression workflow—expression strain, inducer, growth medium, and plasmid—revealed strong convergence toward a relatively restricted set of configurations. Among the 2,388 proteins for which all four parameters were reported (19.6% of the dataset), only 113 distinct configurations were observed. BL21(DE3)-derived expression systems combined with IPTG induction, LB-derived media, and pET-series plasmids formed the single most frequent workflow, accounting for 25.0% of these proteins (*n* = 597); the five and ten most common configurations accounted for 46.0% and 60.6%, respectively. This concentration exceeds what the individual parameter frequencies would predict: given the marginal frequencies of BL21(DE3) (61.6%), IPTG (67.3%), LB (55.9%), and pET vectors (48.1%) within this subset, the leading configuration would be expected in 11.2% of cases if the four choices were made independently, versus the 25.0% observed—a 2.2-fold enrichment. These findings indicate that the convergence observed for individual expression parameters extends to combinations of experimental conditions, with recurrent configurations defining a common recombinant expression infrastructure across diverse proteins and experimental contexts (**Fig. 7**).

**Figure 7.**
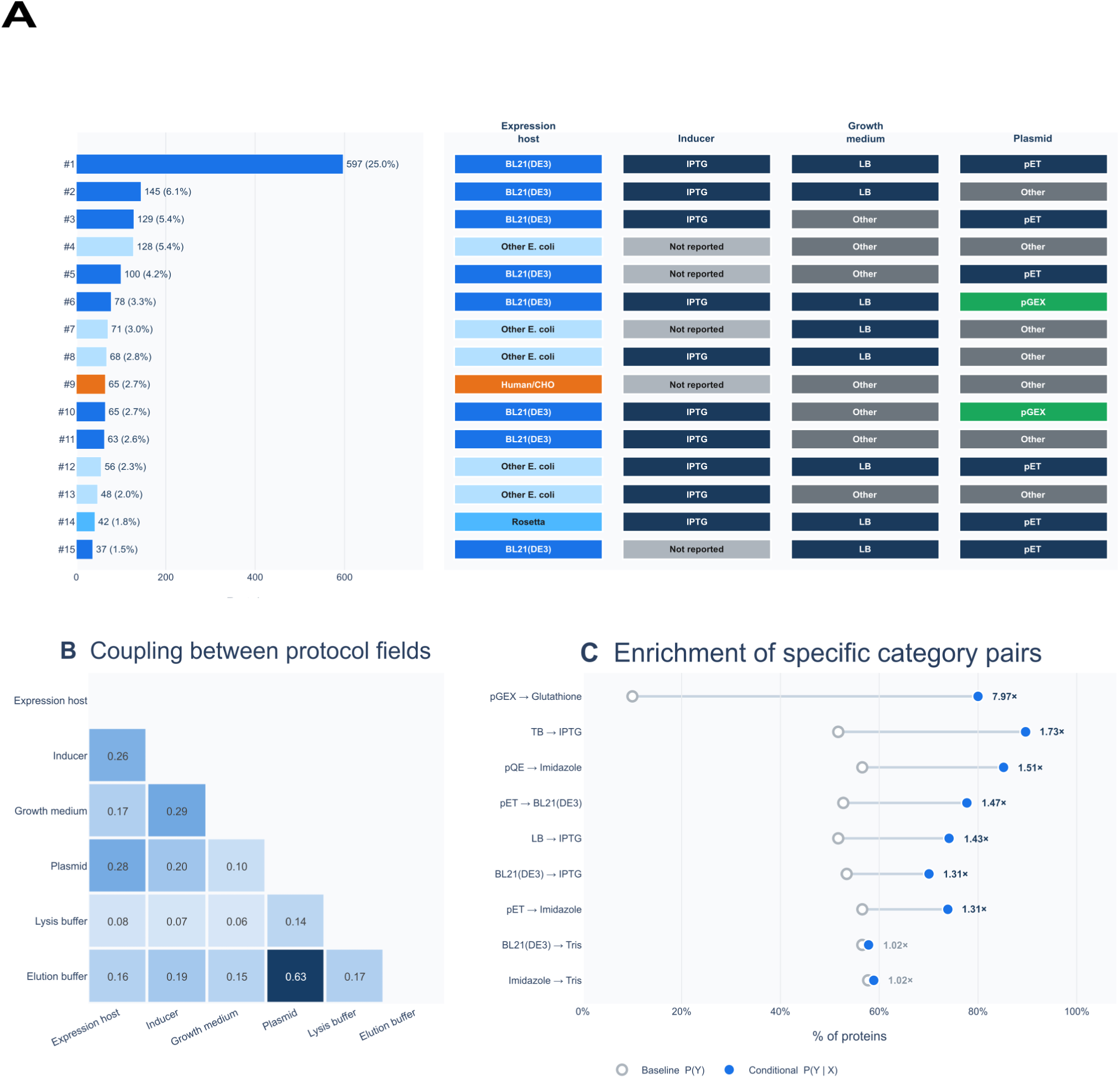
Recurrent purification configurations reveal structured associations among experimental conditions. (A) Most frequently observed recombinant expression configurations, defined by combinations of expression host, inducer, growth medium, and plasmid. Bars indicate the number of proteins associated with each configuration. (**B**) Pairwise associations among purification parameters. Cramér’s V quantifies statistical dependence between categorized protocol fields, with larger values indicating stronger association. (**C**) Enrichment of selected condition pairs, comparing the baseline frequency of each condition with its conditional frequency given the paired condition; values indicate fold enrichment.

Configuration frequencies establish that certain parameter combinations recur, but not which parameters are actually linked. We therefore quantified the pairwise association between protocol fields using Cramér’s *V* (**Fig. 7B**). Coupling was highly uneven across the fifteen field pairs. Plasmid and elution buffer were by far the most strongly associated (*V* = 0.627), 2.19-fold the next strongest pair; inducer and growth medium (*V* = 0.286), expression host and plasmid (*V* = 0.276), and expression host and inducer (*V* = 0.261) formed a second tier.

The strong association between plasmid and elution buffer has a direct biochemical basis and provides a positive control for the co-occurrence analysis: expression vectors can encode affinity tags that determine the compatible affinity-purification and elution chemistry. For example, pGEX vectors encode glutathione S-transferase (GST) fusion tags, which are commonly purified by glutathione-affinity chromatography and eluted with reduced glutathione. Accordingly, although glutathione elution was used in only 10.0% of records reporting both parameters, it was observed in 80.0% of records using a pGEX vector (*n* = 430), corresponding to an 8.0-fold enrichment. Similarly, pMAL vectors encode maltose-binding protein (MBP) fusion tags, which can be purified using amylose-affinity chromatography and eluted with maltose. Consistent with this established workflow, maltose elution occurred in 69.8% of pMAL records compared with 1.0% overall (*n* = 43). Recovery of these established biochemical relationships demonstrates that the reconstructed dataset preserves mechanistically meaningful dependencies between purification parameters.

Enrichments within the expression module were more modest in relative terms because the categories involved are already dominant: pET vectors were paired with BL21(DE3) in 77.8% of cases against a 52.7% baseline (1.5-fold), LB medium with IPTG induction in 74.1% against 51.7% (1.4-fold), and BL21(DE3) with IPTG in 70.1% against 53.4% (1.3-fold). These associations further reflect the reuse of established combinations of expression vectors, hosts, media, and induction systems across recombinant protein production.

Against this background, lysis buffer composition stood out as the one parameter that appears to have the weakest coupling (*V* = 0.065–0.167), indicating that lysis conditions remain comparatively flexible within otherwise recurrent expression and purification workflows.

Together, these results demonstrate that ProtoPure can identify and quantify common practices in protein expression and purification across the scientific literature. Although many of these practices are well established within the experimental community, their prevalence and relationships across the broader literature have remained difficult to quantify systematically. By transforming dispersed protocol descriptions into structured and comparable data, ProtoPure makes this accumulated experimental knowledge accessible at a population scale, enabling established practices to be measured, compared, and placed in context. Efforts such as ProtoPure are therefore important for converting decades of experimental experience embedded in the literature into reusable data that can support systematic analysis, improve access to existing experimental knowledge, and provide a foundation for future data-driven approaches to experimental design.

## DISCUSSION

Protein purification is essential to virtually every experimental protein study; however, knowledge of how to establish effective purification conditions remains largely empirical and fragmented across the literature, making it difficult to systematically reuse existing knowledge or learn generalizable principles from previous experiments.

ProtoPure not only recovers, evaluates, and normalizes protein purification information reported across publications, but also enables the identification of global patterns that emerge throughout the purification process. This shift from individual protocols to population-level analysis provides a framework for examining how experimental decisions are distributed, coupled, and reused across diverse proteins.

Despite analyzing proteins originating from a broad range of organisms and kingdoms of life, we observed a strong tendency for recombinant expression to converge toward Escherichia coli-based systems. This widespread adoption of a common expression platform was accompanied by convergence in several experimental choices, including expression hosts, induction systems, growth conditions, and affinity purification strategies. In contrast, other components of the workflow, particularly buffer composition and downstream purification conditions, remained substantially more heterogeneous and weakly coupled to the surrounding experimental workflow, potentially reflecting the need to accommodate the biochemical properties of individual proteins. Together, these observations suggest that protein purification protocols are organized around a limited set of transferable experimental architectures that combine standardized recombinant expression infrastructure with more adaptable biochemical components.

More broadly, the ProtoPure framework demonstrates that as the scientific literature becomes increasingly machine-readable, experimental workflows themselves can be studied as structured systems at scale. Rather than serving solely as repositories of methodological details, decades of published protocols can be transformed into quantitative datasets that enable the discovery of hidden experimental architectures, recurring design principles, and sources of methodological variability that would be difficult to identify through manual curation alone. Such structured representations provide an opportunity to build upon decades of accumulated experimental knowledge, enabling the development of multimodal representations that integrate protein properties with experimental conditions, as well as specialized language models fine-tuned on protein purification workflows. Ultimately, these approaches could support the development of data-driven frameworks for protein purification and experimental design.

### Limitations of the study

A primary limitation of this study is that ProtoPure reconstructs purification workflows from information explicitly reported in the scientific literature. Consequently, the resulting dataset is influenced by both incomplete reporting and sampling biases in the published record. Many purification parameters are inconsistently documented across publications, and successful purification strategies are likely to be disproportionately represented relative to unsuccessful experiments or conditions that were tested but not reported. Moreover, the current dataset does not systematically associate purification protocols with experimental outcomes such as protein yield, purity, soluble fraction, activity, stability, aggregation, or purification failure. Therefore, the recurrence or enrichment of a particular purification strategy should not necessarily be interpreted as evidence that it is optimal for a given protein. Incorporating quantitative experimental outcomes and unsuccessful purification attempts will be important for moving beyond the descriptive analysis of purification practices toward predictive models capable of recommending experimental conditions.

A second limitation arises from the semantic representation and clustering used to standardize heterogeneous protocol descriptions. The number and composition of the resulting clusters depend on how effectively the embedding model captures similarities between experimentally equivalent conditions while distinguishing biologically meaningful differences. Although the current approach enables large-scale comparison of purification parameters that are reported using highly variable terminology, imperfect representations may either separate equivalent experimental conditions or merge conditions that differ in experimentally relevant ways. Future domain-specific embedding models trained explicitly on biochemical protocols and experimental conditions could provide more biologically informed representations of purification workflows and improve the resolution and robustness of downstream analyses.

## METHODS

### Construction of manually curated protein purification benchmark datasets

To develop and evaluate ProtoPure, we constructed manually curated benchmark datasets for two protein families: azoreductases and S-adenosylmethionine-dependent methyltransferases (SAM-MTases). Azoreductase publications were identified by querying PubMed using the keywords “azoreductases” and “bacteria.” From the retrieved publications, we selected studies reporting protein purification and enzymatic assays, resulting in a benchmark dataset comprising 31 unique publications and 56 protein purification records. SAM-MTase publications were identified from a UniProt-derived table of associated PubMed identifiers (PMIDs). Of 2,332 accessible SAM-MTase publications, 292 were evaluated for the presence of protein purification information; 85 contained explicit purification conditions, whereas 207 were manually classified as lacking purification information. From the purification-positive publications, a subset of 43 publications was randomly selected for manual curation of protein-level purification records. This subset was used to evaluate the accuracy of purification-condition extraction, whereas the full set of publications containing both positive and negative examples was used to evaluate publication-level classification performance.

For both protein families, methods sections were manually extracted from the corresponding publications and used as reference text for evaluating automated methods-section extraction. Protein purification conditions were additionally curated at the protein level according to the predefined 12-field schema shown in Table 1. These manually curated annotations served as ground-truth references for evaluating purification-condition extraction.

The azoreductase dataset was used during pipeline development, with prompts and parameters iteratively adjusted based on performance on this dataset. Because the azoreductase dataset was intentionally restricted to publications reporting experimental purification, it contained only positive examples and was used to evaluate recovery of known purification-positive publications rather than binary classification performance. To evaluate generalization beyond the development dataset, the SAM-MTase dataset was reserved as an independent validation set.

### Classification of Publications According to Presence of Protein Purification Conditions

To ensure that the pipeline accurately identifies publications reporting protein purification conditions, we applied an automated document classification step using LlamaParse. The LLM-based classifier was instructed to produce (i) a binary classification distinguishing protein purification–related from non-protein purification publications and (ii) a concise justification supporting each classification decision. Publications classified as non-protein purification were excluded, while positively classified documents were propagated to subsequent pipeline stages. The prompts used are provided in **Prompt S1**.

Classification performance was evaluated against manually curated ground-truth labels from the azoreductase and SAM-MTase datasets. The azoreductase dataset consisted exclusively of purification-positive publications and was used to evaluate recovery of known positive examples. The SAM-MTase dataset contained both purification-positive and purification-negative publications and was used to evaluate discrimination between the two classes using a confusion matrix and standard classification metrics.

### Extracting Method Sections from Publications

To further simplify the extraction process, we used the OpenAI API to prompt the model to return only the methods section of each paper as plain text. Each document was uploaded to the OpenAI API. Because of the computational cost associated with extracting method sections with multiple GPT models, this step was performed exclusively using GPT-5, which was the most advanced model at the time. The prompt used for this step of the pipeline is shown in **Prompt S2**.

To evaluate extraction accuracy, model-extracted methods sections were compared against manually curated reference sections from the azoreductase and SAM-MTase benchmark datasets using BERTScore F1^23^ employing the SciBERT model^5^, BLEU^29^, cosine similarity, METEOR^30^, and ROUGE-F1^31^ (Fig. S1). BLEU was computed at the unigram level (weights 1, 0, 0, 0) with smoothing; ROUGE-1 F was computed with Porter stemming, and METEOR with WordNet-based stem and synonym matching. Cosine similarity was computed over TF–IDF vectors of word unigrams and bigrams and is reported as a lexical baseline, as implemented in scikit-learn (v1.7.2)^32^.

Metric scores were calculated independently for each publication and subsequently averaged across publications within each benchmark dataset. Error bars represent the standard error of the mean (SEM).

### Extraction of Protein Purification Conditions

To capture the complexity of protein purification workflows, we defined a structured schema encoding the key attributes of the purification process. The schema contains information about the origin of the protein, purification steps, materials and chemicals used during the purification process. The information collected is shown in Table 1. The LLM was instructed to extract and format information consistently across publications.

We evaluated two approaches for providing the methods sections to the LLM, a Retrieval-Augmented Generation approach and a “Prompt insertion” approach, where the methods section extracted in the previous step was manually added. For this stage we evaluated multiple models from OpenAI: GPT-4.1, GPT-5-mini and GPT-5. Similarly, the prompt used for this step is shown in Prompt S3.

The Retrieval-Augmented Generation approach was implemented using the LangChain library^33^ with a Chroma Vector Store^34^ using a chunk size of 600 and chunk overlap of 100. For each publication, retrieval was restricted to chunks belonging exclusively to that document via a metadata filter on the source field. Relevant chunks were retrieved using Maximal Marginal Relevance (MMR), a diversity-aware strategy that balances query relevance with inter-chunk dissimilarity, returning the top 15 candidates.

Prompt insertion was selected for the full-scale pipeline because it showed marginally better overall extraction performance than the RAG-based approach across the benchmark datasets.

Extracted purification conditions were evaluated against the manually curated ground-truth annotations from the azoreductase and SAM-MTase benchmark datasets using the complementary metrics described above.

Because purification conditions can be expressed using semantically equivalent but lexically different descriptions, conventional text-similarity metrics may underestimate extraction accuracy. We therefore complemented these metrics with an LLM-based evaluation using GPT-5-mini. The model was provided with the ProtoPure output and corresponding manually curated ground truth and instructed to assign a similarity score from 0 to 10 according to a predefined rubric, where 10 represented semantic equivalence and 0 represented unrelated or contradictory information. The complete evaluation rubric is provided in Prompt S4.

### Agglomerative Clustering and Standardization by Semantic Similarity

To demonstrate the scalability of the pipeline, we later analyzed publications associated with the proteomes *Mycobacterium tuberculosis (Mtb)*, *Escherichia coli (Ec)* and *Thermus thermophilus (Tt)* as shown in their Swiss-Prot/UniProt tables (UP000001584, UP000000625, UP000000532 respectively). We also employed publications from the protein families Short-Chain Dehydrogenases/Reductases (SDRs), Small GTP-binding domain (SgtPB), and the Heat shock protein 70 family (HSP-70) (InterPro IDs of IPR002347, IPR005225, IPR013126, respectively). The total number of publications analyzed for each dataset is reported in **Fig. 2**.

Publications associated with *Mtb*, *E. coli*, *Tt*, SDRs, SgtPB and HSP-70 were identified using the PMIDs listed in the UniProt tables while ensuring that publications were not duplicated across groups. The corresponding articles were retrieved using the Paperscraper library^35^, which accesses the PubMed Central, Wiley TDM, and Elsevier APIs. The publications that were available on the aforementioned APIs were downloaded and processed through the complete ProtoPure pipeline. The final goal is to retrieve the purification conditions from all the publications of these groups.

Each extracted purification field was independently encoded using a BERT-style sentence encoder to obtain a contextual embedding representing its semantic content. These representations were clustered to identify groups of semantically equivalent purification conditions using the hierarchical community-detection algorithm implemented in the SentenceTransformers library.^36^ For each community, the condition whose embedding was closest to the cluster centroid was selected as the representative condition and assigned as the standardized label for all members of that community.

### Grid Search of Embedding Models and Agglomerative Clustering Parameters

To identify the embedding model and clustering parameters that produced the most coherent semantic clusters, we performed a grid search across 10 BERT-style encoders, five cosine-similarity thresholds, and five minimum cluster sizes. The tested models were: SciBERT^5^, Bioformer^37^, BioBERT Large and BioBERT^38^, SapBERT^39^, BioSimCSE-BioLinkBERT^40^, PubMed-BERT^41^, BIOLORD-2023^42^, all-MiniLM-L6-v2 and all-mpnet-base-v2^43^. The cosine-similarity thresholds used for the grid search were: 0.5, 0.6, 0.7, 0.8 and 0.9 and the minimum cluster sizes were 2, 5, 10, 20, and 30. This resulted in 250 parameter combinations evaluated independently for each purification field.

Each clustering configuration was evaluated according to measurement of how well a sample fits within its assigned cluster compared to neighboring clusters (Silhouette score) and quantification of the average similarity between each cluster and its most similar counterpart (Davies–Bouldin index). The final set of hyperparameters was chosen by maximizing the Silhouette score (higher values indicate better cluster cohesion and separation) and minimizing the Davies–Bouldin index (lower values indicate more compact and well-separated clusters). See **Figs. S3-S13**.

High Silhouette scores and low Davies–Bouldin indices were used to identify well-separated and internally coherent clusters, while the number of clusters was additionally considered to avoid degenerate solutions in which heterogeneous purification conditions were collapsed into a small number of broad communities.

### Protein Purification Analysis with Cluster Analysis

#### Cluster Abundance and Dominance

All abundance analyses were performed on the clustered field values produced by the configuration BioSimCSE-BioLinkBERT-BASE embeddings, minimum community size 2, cosine similarity threshold 0.9. For each purification field independently, records not assigned to any community (cluster identifier *−*1) were discarded, as were clusters whose representative label was a missing-value placeholder (“not mentioned”, “not specified”, “not reported” or “none”), so that all reported statistics describe protocols that were actually specified in the source text. For each remaining cluster *i* we counted the number of records assigned to it, *n_i_*, and its fraction of the field total (Eq. 1), where *K* is the number of clusters recovered for that field. Clusters were ranked by *n_i_*in descending order, and the percentages quoted in the Results for dominant conditions are the *p_i_* of the highest-ranked clusters. Because each record is assigned to exactly one cluster per field, the *p_i_*of a field sum to one and the ranking is a partition of the reported protocols rather than a set of overlapping categories.

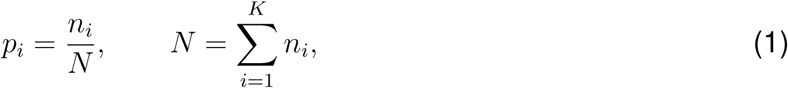

#### Cluster Diversity

Two complementary statistics were computed per field from the same ranked abundance distribution. The first is the raw number of distinct clusters, *K*, which counts how many separable variants of a parameter the corpus contains. The second is the Shannon entropy of the cluster-size distribution and its exponential, the effective number of clusters, with *p_i_* the fraction of records assigned to cluster *i* as defined in Eq. 2.

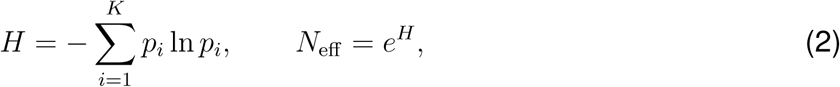

*N*_eff_ is the number of equally abundant clusters that would produce the observed entropy, so it discounts *K* by the skew of the distribution: a field dominated by a few large clusters with a long tail of rare ones has *N*_eff_ *≪ K*, whereas a field whose clusters are evenly populated has *N*_eff_ *≈ K*.

#### Co-occurring Configurations

An expression configuration was defined as the ordered tuple of four parameters: expression host, inducer, growth medium and plasmid. To make the tuples comparable across publications, the cluster assigned to each field was first mapped onto a small set of curated categories by rule — expression hosts into BL21(DE3), Rosetta, C41/C43, other *E. coli*, insect (Sf9), yeast and human/CHO; inducers into IPTG, arabinose, other and not reported; media into LB, TB, 2*×*YT, auto-induction, minimal and other; and plasmids into pET, pGEX, pQE, pMAL and other. Records falling in a noise cluster were treated as unassigned for the field concerned.

The analysis was restricted to complete cases: a protein contributed a configuration only if all four parameters were reported and categorised, with no imputation of missing values.

#### Association and Enrichment Between Purification Parameters

Pairwise association between categorised fields was quantified with Cramér’s *V*. For a pair of fields we built the contingency table of their categories over the proteins for which both were reported (pairwise deletion; the sample size *n* therefore differs between pairs and is given with each coefficient), computed the Pearson *χ*^2^ statistic against the independence expectation *E_ij_*= *R_i_C_j_/n*, and normalised it by its maximum (Eq. 3).

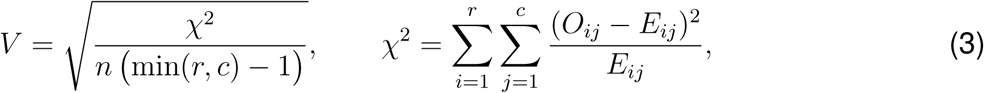

Here *r* and *c* are the numbers of rows and columns of the table and *O_ij_* the observed counts. *V* is symmetric, undirected and bounded in [0, 1], with 0 indicating independence and 1 a deterministic relationship. Because *V* is symmetric and does not indicate which categories drive an association, directional enrichment was quantified separately. For a category *a* of field *X* and a category *b* of field *Y*, the conditional frequency is the proportion of records with *X* = *a* that also have *Y* = *b*, and the baseline is the marginal frequency *P* (*Y* = *b*) computed on the same set of complete pairs, so that the two are directly comparable (Eq. 4).

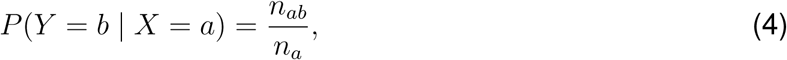

Enrichment was expressed as the ratio of the two, reported on a log_2_ scale, on which zero denotes a conditional frequency equal to the baseline, positive values enrichment and negative values depletion (Eq. 5).

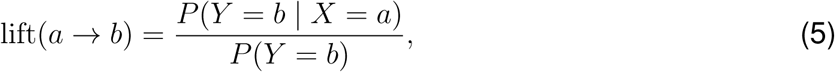

## RESOURCE AVAILABILITY

### Lead contact

Requests for further information and resources should be directed to and will be fulfilled by the lead contact Prof. Adrian Jinich or first author Ricardo Almada Monter.

### Materials availability

This study did not generate new materials.

### Data and code availability

- Extracted protein purification conditions are publicly available on the ProtoPure official website at https://huggingface.co/spaces/richiam/ProtoPure.
- Source code and analysis scripts are available on GitHub at github.com/jinichlab/llm extractor.
- Additional information required to reanalyze the data reported in this paper is available from the lead contact upon request.

## ACKNOWLEDGMENTS

This work was supported by HHMI Hanna Gray Fellowship and NIH FIRST Program Fellowship (U54CA272220). The authors would like to thank the participants of the ENLACE 2024 summer program, Olivia A. Graeve and administrators of such programs for their support in the initial collection of data for this project. The authors also would like to thank the UCSD Physics Computing Facility for its support in the maintenance of computational resources.

## AUTHOR CONTRIBUTIONS

Conceptualization: R.A.-M., A.J., and J.M.-L.; Methodology: R.A.-M., A.J., and J.M.-L.; Software: R.A.-M. and J.M.-L.; Validation: R.A.-M. and J.M.-L.; Formal analysis: R.A.-M.; Investigation: R.A.-M. and J.M.-L.; Data curation: R.A.-M., E.G., J.T.-S.A., A. Spencer, A. Siordia, P.C., S.L.-A., J.E.-H., N.R., S.M.V., A.U., M.S., and A.J; Visualization: R.A.-M. and J.M.-L.; Writing – original draft: R.A.-M. and A.J.; Writing – review & editing: R.A.-M. and A.J; Supervision: A.J.; Funding acquisition: A.J.

## DECLARATION OF GENERATIVE AI AND AI-ASSISTED TECHNOLOGIES

During the preparation of this work, the authors used ChatGPT to assist with manuscript editing and language refinement. After using this tool, the authors reviewed and edited the content as needed and take responsibility for the content of the publication.

## SUPPLEMENTAL INFORMATION INDEX

Prompt S1. Classification Prompt

Prompt S2. Methods Extraction Prompt

Prompt S3. Purification Extraction Prompt

Prompt S4. LLM as a Judge of Purification Protocol Reconstruction

Figure S1. Metric Evaluation on Method Section Extraction for Azoreductases and SAMMTases

Figure S2. BERTScore values for the evaluation of the retrieved information in the Azoreductases (Top figure) and the SAM-MTases (bottom)

Figure S3. Grid Search Parameters Summary

Figure S4. Internal clustering metrics across the parameter grid — SciBERT.

Figure S5. Internal clustering metrics across the parameter grid — Bioformer.

Figure S6. Internal clustering metrics across the parameter grid — BioBERT Large.

Figure S7. Internal clustering metrics across the parameter grid — BioBERT.

Figure S8. Internal clustering metrics across the parameter grid — SapBERT.

Figure S9. Internal clustering metrics across the parameter grid — BioSimCSE-BioLinkBERT.

Figure S10. Internal clustering metrics across the parameter grid — PubMedBERT.

Figure S11. Internal clustering metrics across the parameter grid — BioLORD-2023.

Figure S12. Internal clustering metrics across the parameter grid — all-MiniLM-L6-v2.

Figure S13. Internal clustering metrics across the parameter grid — all-mpnet-base-v2.

