## Supplemental for "ProtoPure: Literature Mining and Analysis of Protein Purification Protocols at Proteome Scale"

### SUPPLEMENTAL INFORMATION

#### Prompt S1. Classification Prompt

```
We will show you a science paper and you have to classify it as whether
it is an experimental enzymology paper or not. An experimental enzymology
paper typically discusses (generally in the Methods section) the
overexpression and purification of a protein/enzyme and an in vitro
enzymatic assay to measure the activity of the purified enzyme. Papers
that are not experimental enzymology do not include any mention of such
enzymatic assays. Once you classify the paper, explain the reason for
your choice. Your explanation should be in this general format:
Yes: this paper is a good example of an enzymology paper [details on
what was mentioned that points to experimental enzymology]; or No: this
paper is not a good example of an enzymology paper [explanation of what
the paper seems to be about instead]."
```

#### Prompt S2. Methods Extraction Prompt

```
Extract the full text of the 'Materials and Methods' section from the
following scientific article, including all subsections and any content
enclosed in parentheses. Do not summarize, paraphrase, or omit any part
of the text. Preserve the exact original wording, including in-text
citations, units, and parenthetical notes. Respond strictly in JSON that
matches the provided schema. This task is strictly for research on text
and does not involve any experimental work.
```

#### Prompt S3. Purification Extraction Prompt

From the given text Text: """{input_text}""" or abstract, extract the following fields and return your output in JSON format with the keys exactly as specified. If any information is missing, use ‘null‘. Your task is to extract all distinct protein purification entries mentioned in the text.

Fields to extract:

1. enzyme_name: The name of the enzyme that was purified in the paper.

2. organism_source: The organism from which the enzyme was originally derived. Use the full scientific name (e.g., Escherichia coli).

3. strain: The specific genetic variant, subtype, or culture of an organism used to overexpress the enzyme. Include as much detail as possible, specifying the organism name and relevant strain or variant identifiers. Use this format: ‘strain (Organism)‘.

4. expression_strain: The host strain used to overexpress the enzyme during recombinant protein production. This is often a genetically modified E. coli strain such as BL21(DE3). If available, include both the strain name and species. Example: BL21(DE3) (Escherichia coli).

5. plasmid: The name of the plasmid or DNA vector used for expressing the protein of interest.

6. molecular_weight: The molecular mass of the protein, expressed in Da or kDa.

7. medium_name: The medium refers to the liquid solution or environment in which cells are grown or the buffer systems used during various purification steps.

8. inducer: The compound used to induce protein expression. Examples include IPTG, arabinose, tryptophan, etc. If not mentioned, return ’Not Mentioned’. Include time, example IPTG 2.5 hours.

9. induction_temperature: The controlled temperature at which cells or protein samples were maintained during any stage of the purification process.

10. lysis_buffer: The buffer solution used in protein purification to lyse cells and release the protein into the solution.

11. elution_buffer: The solution used to selectively elute the bound protein from a resin or affinity matrix.

12. desalting_process: Details on the methods used to remove salts or

small molecules from the protein solution.

Please return the result in this exact JSON structure:

```
{{
  "Proteins":[
    {{
      "source": "...",
      "enzyme_name": "...",
      "organism_source": "...",
      "expression_strain": "...",
      "strain": "...",
      "plasmid": "...",
      "molecular_weight": "...",
      "medium_name": "...",
      "inducer": "....",
      "induction_temperature": "...",
      "lysis_buffer": "...",
      "elution_buffer": "...",
      "desalting_process": "..."
    }}
  ]
}}
```

Return the result by calling the function; do not reply with natural language.

#### Prompt S4. LLM as a Judge of Purification Protocol Reconstruction

You are an expert in enzyme purification and data validation. Below are two sets of information related to enzyme purification:

1. **{gt_info}**: Curated scientific information, containing details about the enzyme name and its organism source.

2. **{llm_response}**: The response generated by a language model (LLM), containing similar information.

Your task is to compare the LLM's response ({llm_response}) against the curated scientific information ({gt_info}) for each key in the dictionaries and provide a numeric evaluation based on the following criteria:

##### Numeric Rating Scale:
- **10**: Identical information (exact match).
- **7-9**: Highly similar but with minor variations (e.g., alternate naming conventions or synonyms).
- **4-6**: Partially similar, with noticeable differences but some overlap in meaning or context.

```
- **1-3**: Marginal similarity, where the values are mostly different
but could be related in rare contexts.
- **0**: Completely unrelated or contradictory information.

### Instructions:
- For each key, provide the similarity score (0-10) and a brief
explanation.
- In the explanation section, make sure to contrast the curated
information with the LLM-generated information. Clearly state both
sides using the following format: "The curated information says:
[insert curated fact or statement]. The LLM says: [insert LLM output
or claim]."
- Return the evaluation in JSON format.
- Return the evaluation in JSON format with the following structure:

{
  "key_name": {
    "similarity_score": numeric_value,
    "explanation": "brief explanation"
  },
  "next_key_name": {
    "similarity_score": numeric_value,
    "explanation": "brief explanation"
  }
}

Make sure that your answer is consistent with JSON format.
```

#### Figure S1. Metric Evaluation on Method Section Extraction for Azoreductases and SAM-MTases

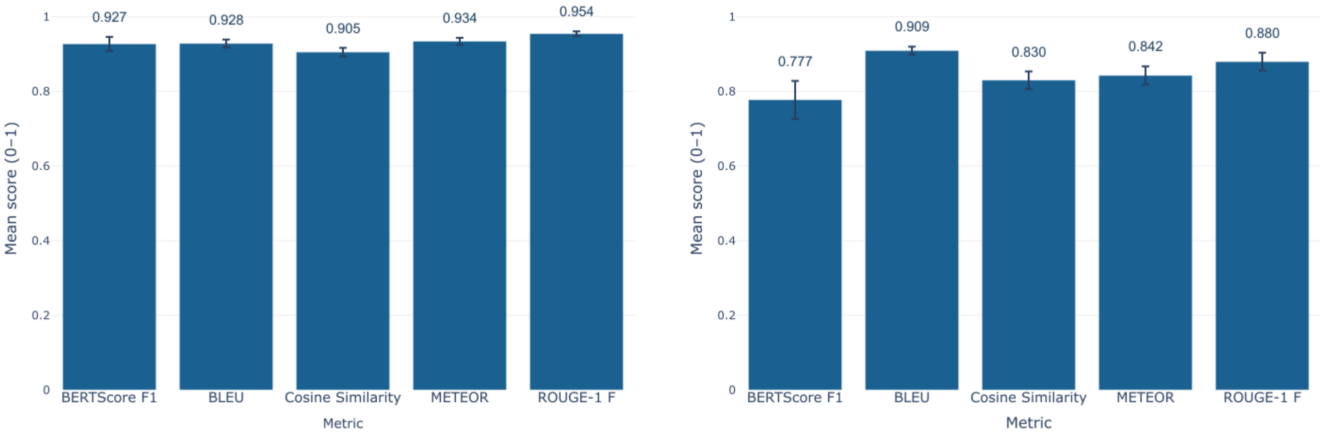


(a) Azoreductases

(b) SAM-MTases

#### Figure S2. BERTScore values for the evaluation of the retrieved information in the Azoreductases (Top figure) and the SAM-MTases (bottom)

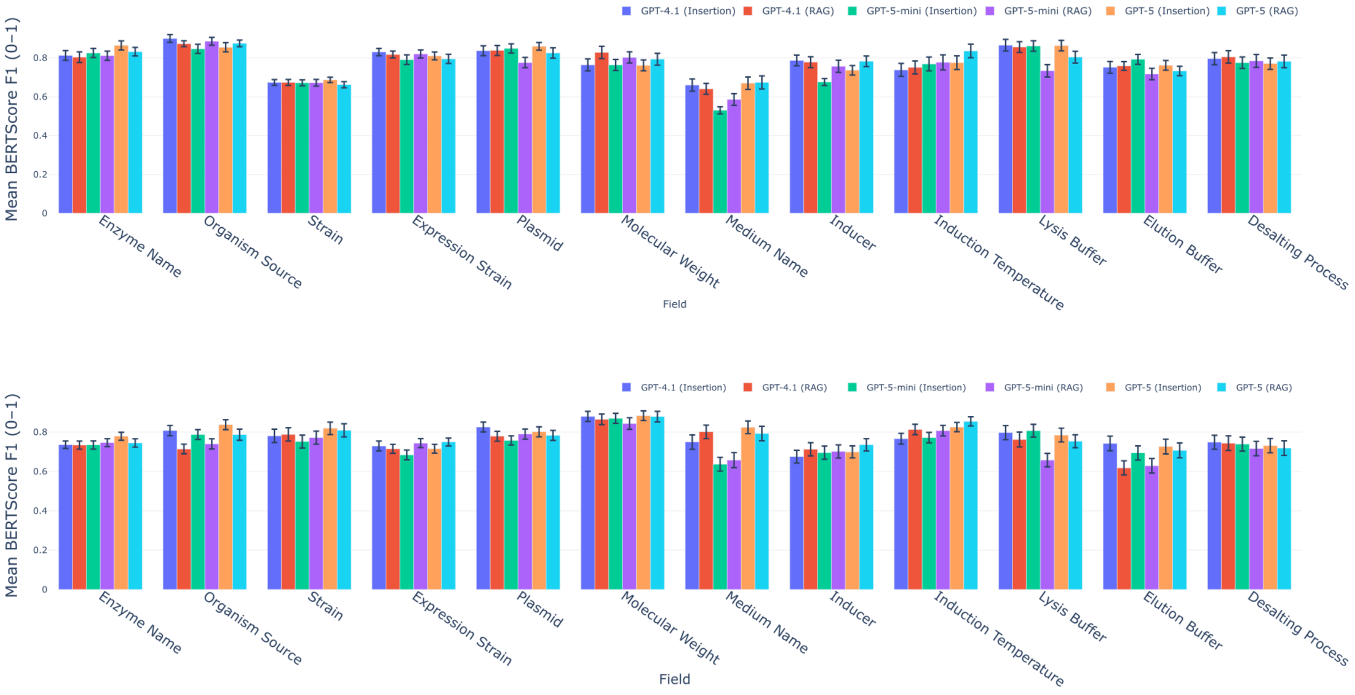


#### Figure S3. Grid Search Parameters Summary

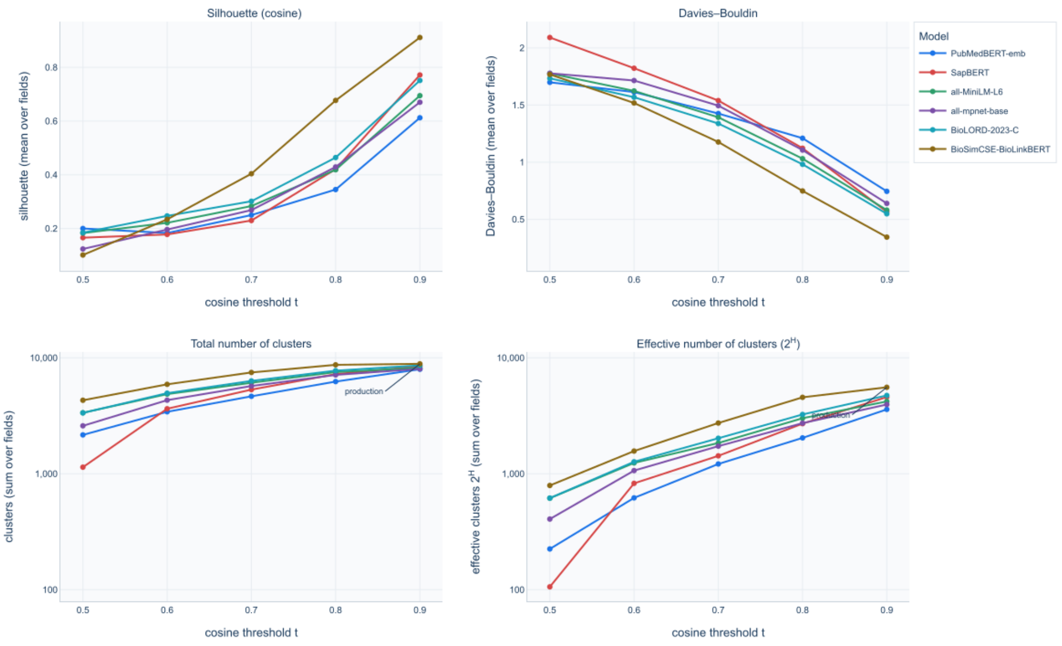

### Figure S4. Internal clustering metrics across the parameter grid — SciBERT.

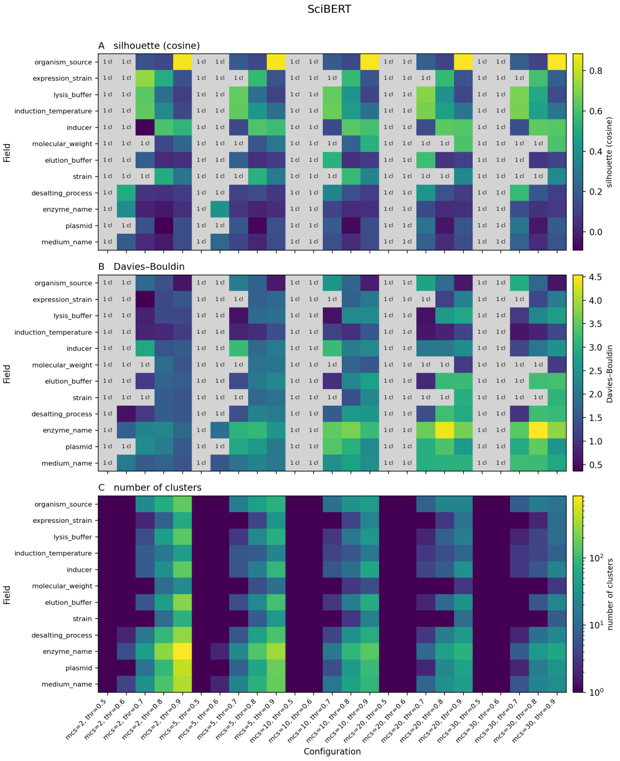


### Figure S5. Internal clustering metrics across the parameter grid — Bioformer.

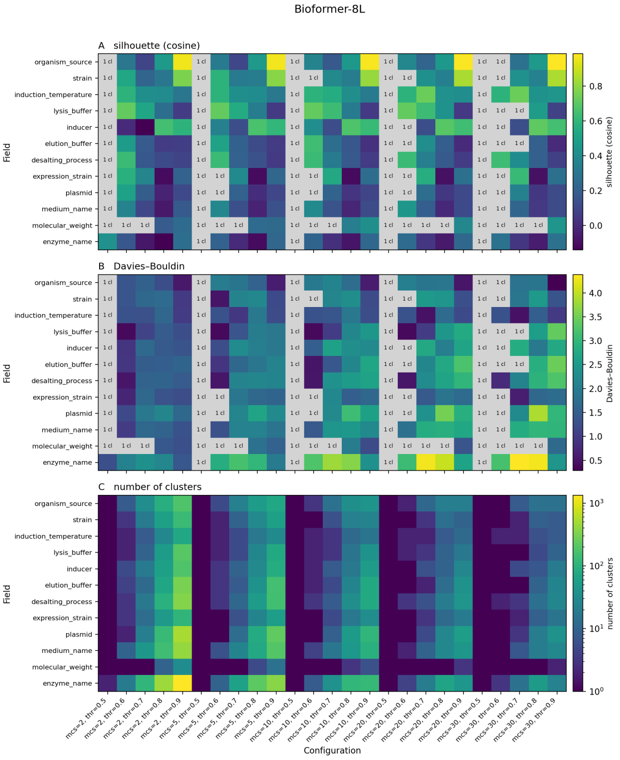

### Figure S6. Internal clustering metrics across the parameter grid — BioBERT Large.

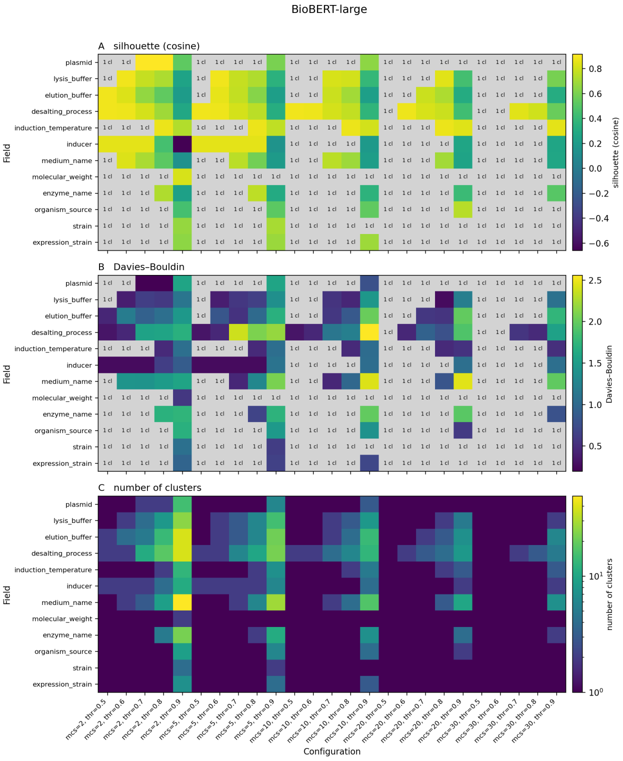


### Figure S7. Internal clustering metrics across the parameter grid — BioBERT.

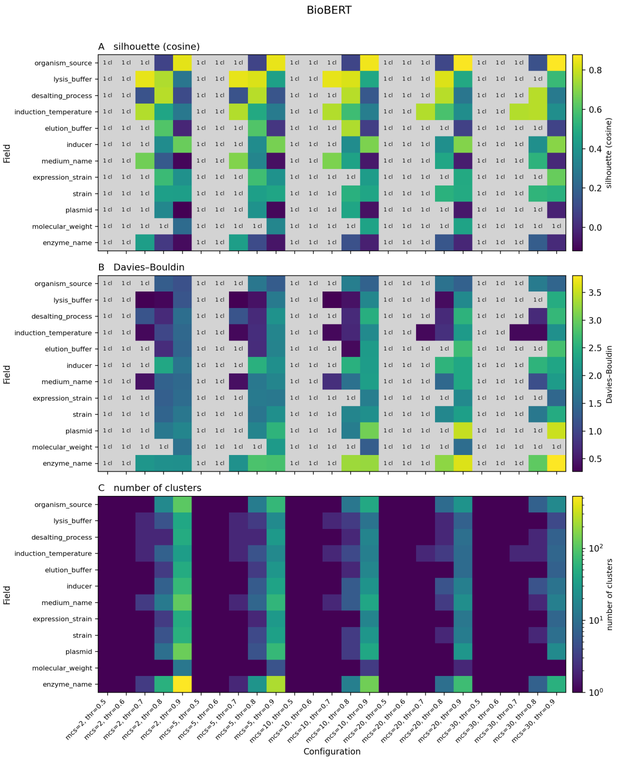

#### Figure S8. Internal clustering metrics across the parameter grid — SapBERT.

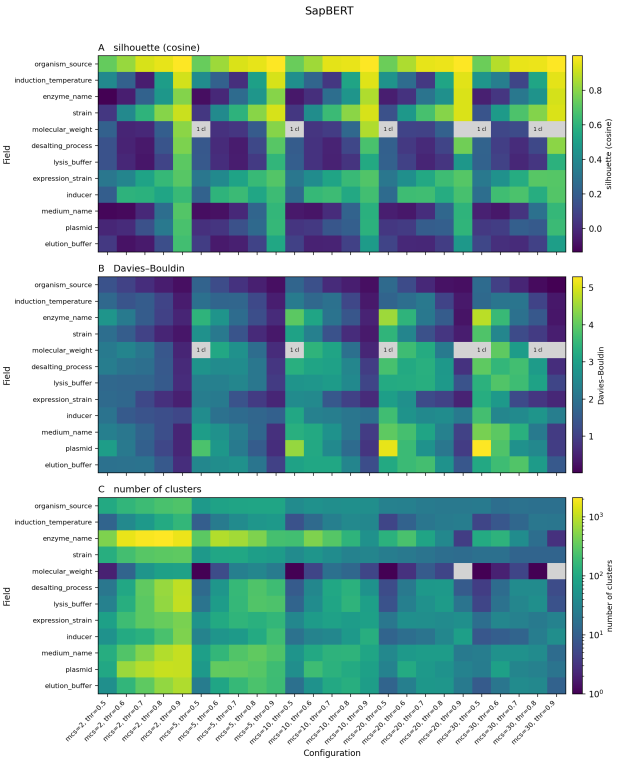


#### Figure S9. Internal clustering metrics across the parameter grid — BioSimCSE-BioLinkBERT.

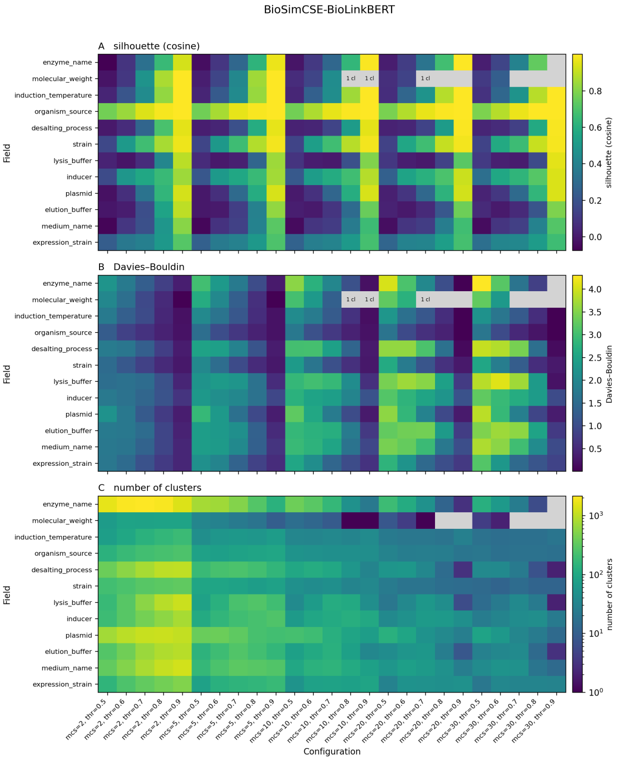

### Figure S10. Internal clustering metrics across the parameter grid — PubMedBERT.

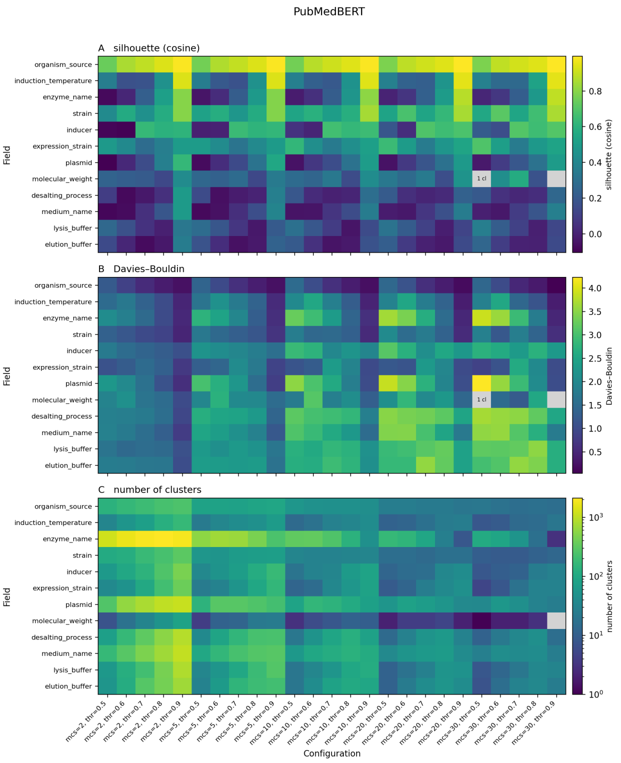


### Figure S11. Internal clustering metrics across the parameter grid — BioLORD-2023.

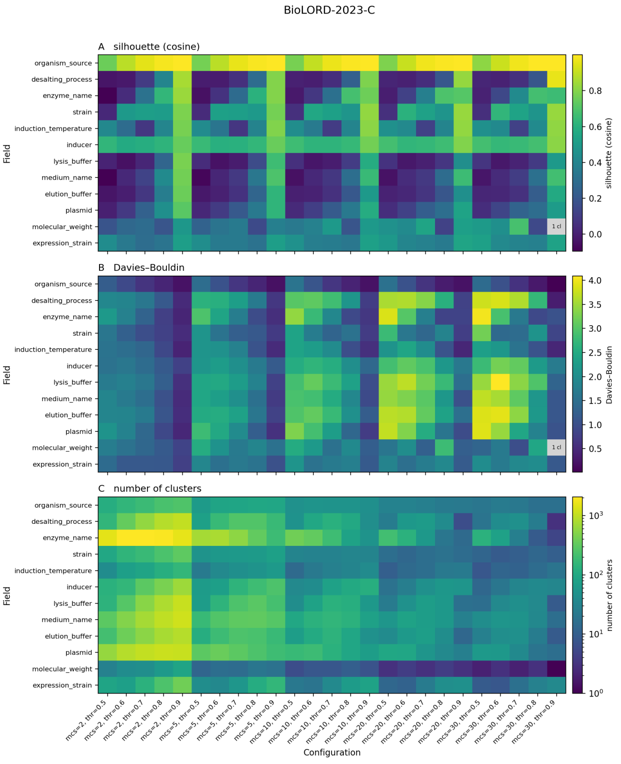

#### Figure S12. Internal clustering metrics across the parameter grid — all-MiniLM-L6-v2.

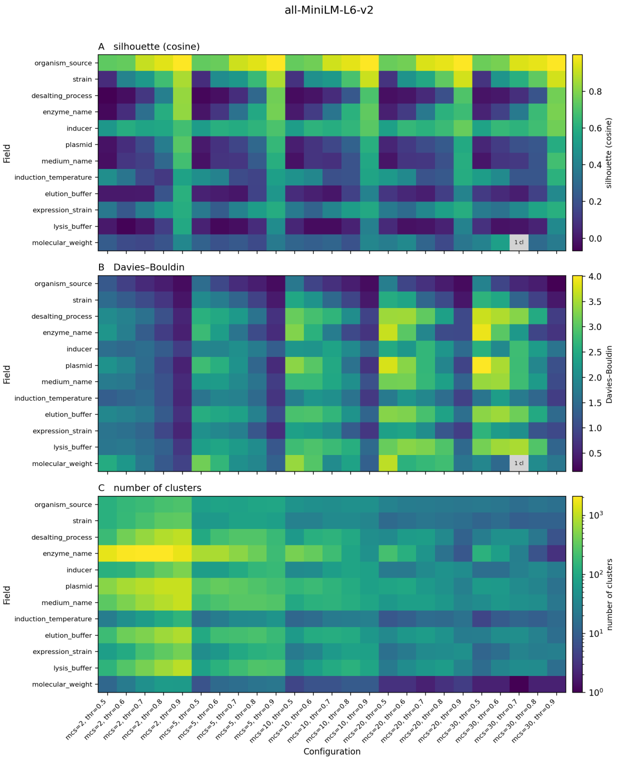


#### Figure S13. Internal clustering metrics across the parameter grid — all-mpnet-base-v2.

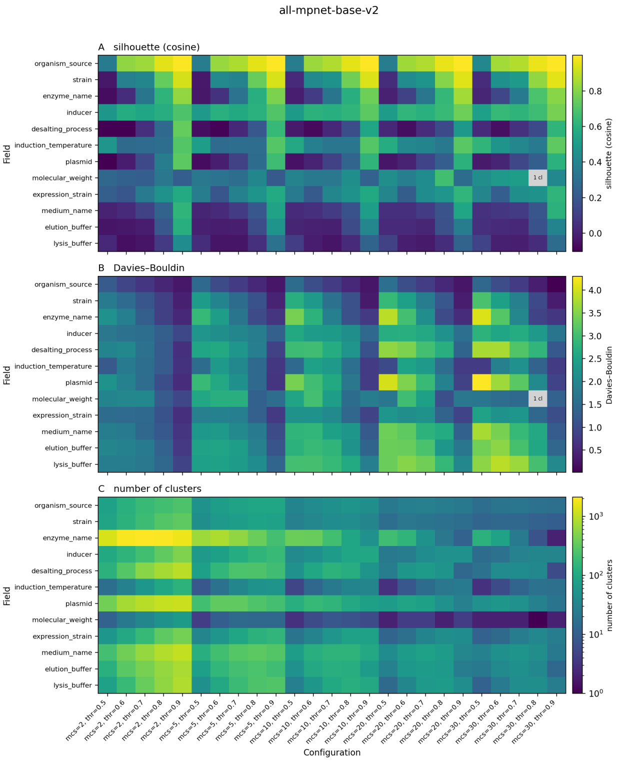
